# NAD(P)H-FLIM reveals spatial and temporal epithelial metabolic heterogeneity in live intestinal organoids

**DOI:** 10.64898/2026.11.25.769250

**Authors:** Hang Zhou, Irina A. Okkelman, Li Lei, Amber De Kinder, Lars Vereecke, Bart N. Lambrecht, Kees Jalink, Ruslan I. Dmitriev

## Abstract

Metabolism contributes globally to energy production at cell and tissue levels, but its cell-specific metabolic states also support tissue function and architecture. Conventionally used bulk metabolic flux assays however provide only the average readouts across cell populations, while fixed-sample and destructive spatial analysis methods result in endpoint. NAD(P)H fluorescence lifetime imaging microscopy (FLIM) enables label-free, spatially resolved imaging of the individual cells and their responses to metabolic perturbations, directly probing dynamics of catabolic and anabolic metabolic pathways. The application of FLIM methodology to the intact living three-dimensional tissues such as stem cell-derived organoids, linking the cellular NAD(P)H dynamics to cell identity and mitochondrial function remains challenging. Here, we combined NAD(P)H-FLIM with lineage reporters, fluorescent probes and pharmacological perturbations in live mouse small intestinal organoids. Optimized acquisition and guided differentiation revealed drastic effects of the culture growth medium and ‘crypt-villus’ axis on the intercellular gradient of NAD(P)H and mitochondrial polarization. Further, guided organoids differentiation and marker-based analyses identified enterocytes as the principal population with highest NAD(P)H intensity and long lifetimes. Enterocyte-confined ‘high’ NAD(P)H correlated with higher cell-specific ATP, contrasting with higher crypt/stem cell niche-associated mitochondrial membrane potential (ΔΨm) that persisted after apical efflux inhibition. Compared to the crypt-based cells, NAD(P)H in villus mitochondria-specific and high ATP enterocyte regions was also more sensitive to oligomycin, suggesting OxPhos as their main energy production mechanism. These findings indicate greater proton-gradient utilization for ATP synthesis potentially contributing to the lower ΔΨm in enterocytes, whereas higher ROS levels were revealed in crypt. Altogether, integrating epithelial lineage assignment with complementary detection of ΔΨm, ROS and ATP provide better insight in interpretation of NAD(P)H-FLIM applied in a broader context of regional cell metabolic heterogeneity, supporting future epithelial dynamic metabolic studies visualizing intestinal regeneration and disease progression.

## Introduction

The small intestinal epithelium is a metabolically active tissue that fulfills a plethora of physiological functions, including rapid renewal of the epithelial cell monolayer, forming the primary barrier against microbiota and toxins, acting as an interface for nutrient absorption and serving as a “hub” for inter-tissue/organ signaling pathways, such as enteric-neural interactions and inflammation^1^. The self-renewal process in the mammalian intestinal tract requires only several days^1^, and involves cell signaling, transcriptional response and an immense metabolic undertaking. To recapitulate this metabolic niche in vitro, an intestinal organoid model, first described by Sato and Clevers^2^, reconstructed a crypt–villus epithelial structure by embedding isolated stem cells into Matrigel hydrogel and supplementation with EGF, Noggin, and R-spondin. Crucially, this organoid model recapitulates major functional features of the intestinal epithelium: it develops budding structures representing the intestinal crypts (stem and Paneth cells) and an internal lumen region lined by differentiated cells (enterocytes, goblet, enteroendocrine^3,4^, and Tuft cells^5^) belonging to the villus domain. Such ‘organized’ cellular heterogeneity and cytoarchitecture enabled intestinal organoids as a promising in vitro model for developmental, biomedical and tissue engineering research. However, even 15 years since their introduction, studying organoid models’ metabolism is still a profound challenge, mostly relying on the use of ‘conventional’ metabolic analysis methods such as bulk extracellular Flux (XF), Fluorescence-Activated Cell Sorting (FACS), mass spectrometry imaging and other end-point methods^6–8^ leading to the loss of crypt-villus spatial or temporal “4D” metabolic complexity^7–9^.

Nicotinamide Adenine Dinucleotide (NADH) and its phosphorylated form (NADPH) are central electron carriers in cell metabolism across all the kingdoms of life^10,11^. NADH is the primary donor of electrons to the mitochondrial Electron Transport Chain (ETC), driving the Oxidative Phosphorylation (OxPhos) process. NADPH is the essential reducing equivalent for anabolic biosynthesis (lipid/nucleotide synthesis) and neutralization of Reactive Oxygen Species (ROS). Critically, NADH and NADPH (collectively referred to as NAD(P)H due to their overlapping emission spectra) are autofluorescent when excited by UV (330-360 nm) or two-photon light (typically 740-760 nm)^10^. Two-photon fluorescence lifetime imaging microscopy (FLIM), also frequently referred to as ‘Optical Metabolic Imaging’, allows for direct, label-free visualization of the NAD(P)H in the cell metabolic pool, separated from other autofluorescent pigments^7,11^. Unlike intensity-based imaging, FLIM calculates fluorescence lifetime (τ), which is largely independent from the fluorophore concentration and image acquisition conditions, but sensitive to the dye (micro)environment due to binding of proteins (e.g. at the ETC), protonation (pH), self-quenching, temperature and viscosity^7^. FLIM effectively distinguishes between the free and protein-bound NAD(P)H forms^12^. The π-π interaction of nicotinamide and adenine rings in free NAD(P)H solution self-quenches the fluorescence resulting in its shortest lifetime (τ_(free)_ = 0.4 ns)^13,14^, typically associated with glycolysis. Upon enzyme binding, the cofactor unfolds, reducing quenching and significantly extending the lifetime to 1.9–5.7 ns^14,15^. Consequently, a shift toward the longer lifetimes is frequently interpreted as an increase in the ‘bound’ fraction, reflecting higher OxPhos activity. Recent research highlights that the ‘classical’ Glycolysis/OxPhos interpretation model requires some corrections: thus, Blacker and co-workers identified that ‘bound NAD(P)H’ is not present in a homogeneous state; NADH and NADPH exhibit distinct lifetimes when bound (1.5 ns vs. 4.4 ns, respectively)^12^ and can reflect multiple binding configurations^16^. Some enzyme-bound states can also exhibit lifetimes comparable to or shorter than free NAD(P)H. Therefore, increased intensity or longer lifetime cannot, by itself, inform on greater OxPhos or ATP production^17^, requiring complementary measurements and defined perturbation conditions to support NAD(P)H-FLIM biological interpretation. A number of complementary approaches have been therefore proposed, including measurement of additional autofluorescent co-factors (FAD, FMN or other spectrally overlapping pigments)^18^, fluorescence anisotropy^16,19^, transcriptome cell specific profiling of NAD(P)H-binding enzymes^20^ or combining with imaging of molecular oxygen (O_2_)^8,21^.

Recently, using organoid models, it has been revealed that the intestinal epithelial cells display distinct metabolic requirements for anabolic biosynthesis and redox homeostasis. Groups of Burgering and Rodríguez-Colman reported that Lgr5^+^ intestinal stem cells (ISCs) exhibit higher mitochondrial OxPhos activity compared to their neighboring Paneth cells^22,23^. Chaves-Perez and co-workers demonstrated that absorptive enterocytes upregulate TCA cycle enzymes (OGDH) for higher OxPhos, while secretory lineages downregulate these pathways^24^. Our team has also shown that stem and differentiated cells in organoids display strong oxidative metabolism, resulting in cellular O_2_ gradients and dynamically responding to the nutrient and growth medium composition^8,25^. Consequently, this leads to the hypothesis that the metabolic landscape of the living intestinal epithelium, reproduced in organoids, is not a simple gradient but a dynamic mosaic of distinct cell-specific bioenergetic states.

Answering the questions on which epithelial lineages account for spatial differences in NAD(P)H metabolism and how are they connected to the energy production at the mitochondria, requires resolving of the cell lineage identity within the living organoid model. Such research endeavor would require comparing cell lineage responses both in specific regions and within the crypt-villus axis as a whole. Therefore, here we investigated these relationships in live mouse small intestinal organoids by combining the NAD(P)H-FLIM method with cell lineage identification and complementary functional imaging dyes. An opposing spatial distribution of enterocyte-associated NAD(P)H autofluorescence and crypt-confined mitochondrial membrane potential provided the central observation motivating our study. We optimized photon collection for single-cell lifetime analysis and used directed differentiation, immunofluorescence and transgenic reporters to identify the cellular origin of NAD(P)H signals. We then examined mitochondrial polarization while accounting for apical dye efflux, compared ATP-sensitive and oxidation-sensitive fluorescence across epithelial regions, and measured cell-dependent NAD(P)H responses upon inhibition of the electron transport, ATP synthase and selected routes of substrate utilization. The presented integrated approach highlights how cell lineage identity and mitochondrial function are linked to endogenous fluorescence, providing a foundation for interpreting dynamic 4D metabolic heterogeneity in the epithelial 3D tissue models beyond regional intensity and lifetime gradients alone.

## Results

### Optimization of NAD(P)H-FLIM acquisition and processing enables single-cell metabolic analysis in live intestinal organoids

While NAD(P)H-FLIM has been already proposed for imaging 3D and in vivo tissues, it has rarely been applied to live high-resolution experiments with living organoid tissues that contain multiple cell types^8,26–31^. Notably, reliable NAD(P)H-FLIM requires sufficient photons for precise lifetime measurements while also limiting phototoxicity, which can influence cellular metabolism. We therefore optimized acquisition settings to balance signal quality, spatial resolution and acquisition speed with minimal light exposure, supporting subsequent biological comparisons. We used GSLab^32^, an open-source phasor analysis software package, to quantify intestinal organoid NAD(P)H intensity and lifetime. Background pixels and luminal autofluorescence were removed, followed by the export of organoid pixel-based NAD(P)H signals as CSV files (Fig. S1A). Using this workflow, we first screened eight two-photon FLIM acquisition settings (Supplementary Table 6) at the organoid level using 740 nm excitation, 410–490 nm emission collection and zoom 1 (Fig. 1A–D). These settings produced a photon-count range from 18.03 to 81.58 photons/pixel. For downstream selection, the settings were grouped into three practical photon count groups: low (18.03–20.05 photons/pixel), intermediate (29.82–40.00 photons/pixel) and high (74.76–81.58 photons/pixel) photon counts. Across these groups, NAD(P)H τ(φ) modestly decreased with increasing photon count, from 1.876 ns in the low-photon group to 1.789 ns in the high-photon group, whereas τ(m) remained comparatively stable. In addition, effect of zoom factor was also tested (Fig. S1 B,C): increasing zoom from 1 to 3 caused slight changes in NAD(P)H photons/pixel, τ(φ), and τ(m), indicating that zoom had limited influence on organoid-level NAD(P)H-FLIM readouts.

**Figure 1.**
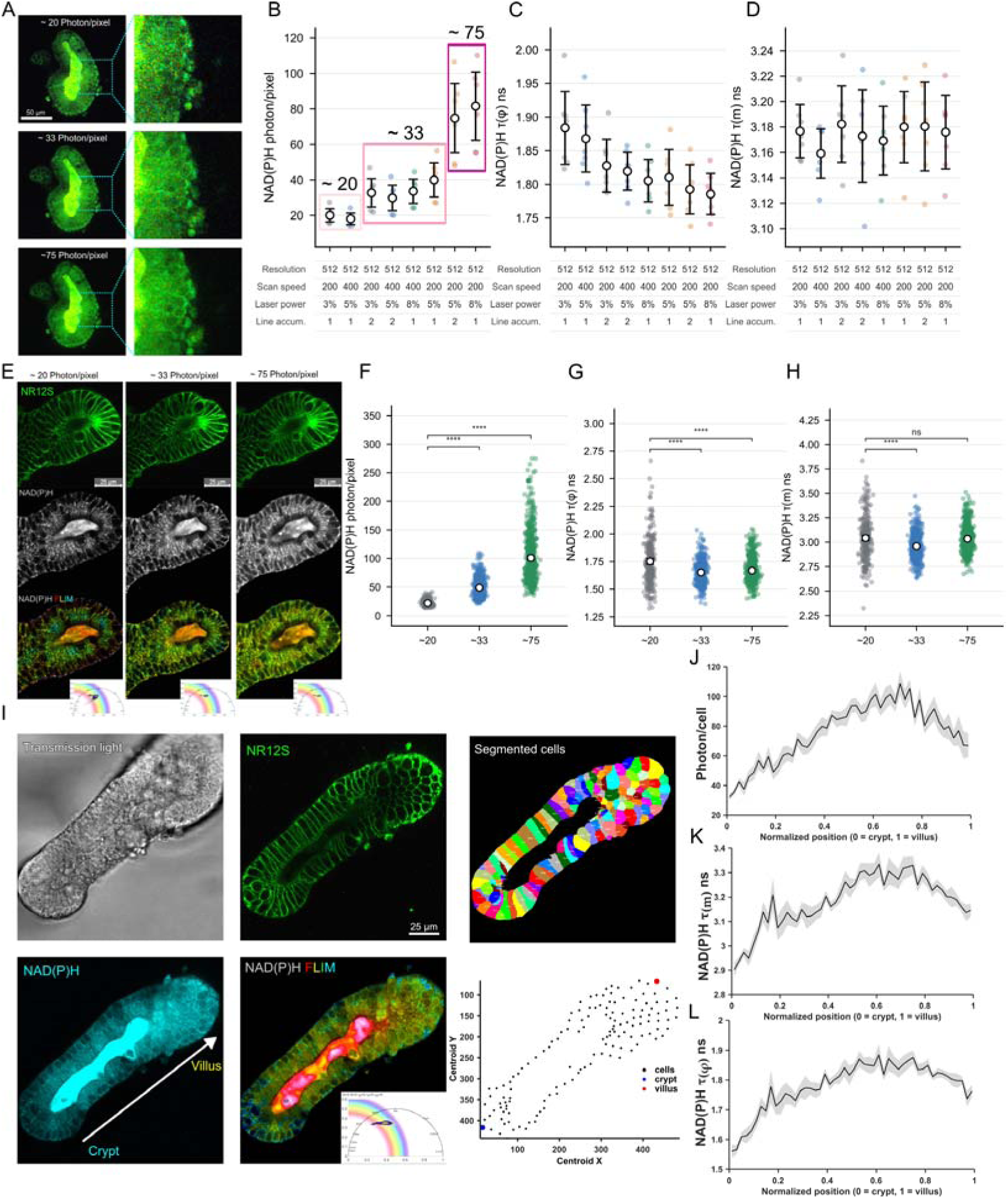
Optimization of two-photon NAD(P)H-FLIM acquisition settings enables sufficient photon collection for visualizing crypt-villus single cell NAD(P)H heterogeneity in small intestinal organoids. A: Representative FLIM images of NAD(P)H in intestinal organoid made to reach average ∼ 20 photon/pixel (Low photon counts), ∼ 33 photon/pixel (Intermediate photon counts), and ∼ 75 photon/pixel (High photon counts). Magnified views are shown on the right. Scale bar: 50 _μ_m. B-D: Imaging settings to gain ∼20, ∼33 and ∼75 photon/pixel for NAD(P)H intensity, with corresponding average NAD(P)H _τ_(_φ_) and _τ_(m) (all imaging settings were done with Exc. 740 nm, Em. 410-490, zoom 1, 7 samples for each group, settings are displayed as Resolution, Scan speed, Laser percentage, Line accumulation, details in Supplementary Table 6). E: NR12S cell membrane staining (intensity) and NAD(P)H (intensity and FLIM) in intestinal organoids at three photon/pixel levels. Phasor plot displays NAD(P)H-FLIM images (rainbow color gradient) where hue represents _τ_(m) magnitude on phasor plot. Scale bar: 25 _μ_m. F-H: Single cell average photons with corresponding average NAD(P)H _τ_(_φ_) and _τ_(m). I: Single cell NAD(P)H intensity and lifetime values across the crypt-villus axis: transmission light, NR12S fluorescence, cells mask from NR12S, NAD(P)H intensity (Cyan), FLIM (false-color gradient from phasor plot), and X, Y centroid of cells. The scale bar in the NR12S image represents 25 µm. Dots (blue for crypt, red for villus) defining cell positions are obtained by applying cell masks to GSLab, along with mean photons and lifetime data of cells. J-L: Line profile across the crypt to villus single cell NAD(P)H photons, _τ_(m) and _τ_(_φ_) area (normalized from 0 to 1 by customized Matlab code, 0 = crypt side, 1 = villus side, 5 samples). Quantitative summaries and existing comparisons are provided in Supplementary Table 8.

We next asked whether these settings could support single-cell FLIM analysis. To achieve this, we first applied NR12S, a Nile Red modified lipid membrane live imaging dye^33^, to reliably label cell boundaries within the organoid epithelium for generating single-cell masks. We then extracted NAD(P)H-FLIM pixels information of individual cells by applying cell masks in GSLab. With the chosen one acquisition setting for each photon-count group, valid single-cell ROIs showed 21.83, 48.91, and 101.00 photons/pixel, respectively. Obtained τ(φ) were 1.752, 1.652, and 1.669 ns, and τ(m) were in the range of 3.042, 2.960, and 3.035 ns (Fig. 1E–H). ROI validity increased with photon count (Fig. S1D). Based on 520, 572, and 622 analyzed cells under low, intermediate and high photons/pixel imaging conditions, valid τ(φ) ROIs increased from 51.15% to 96.51% and 100%, while τ(m) ROIs from 71.15% to 99.13% and 100%. These results indicated that higher photon-count acquisition improved valid single-cell ROI and reduced variability in lifetime calculation, supporting intermediate-to-high photon-count settings for robust single-cell NAD(P)H-FLIM analysis. We next asked whether low-photon single-cell analysis could be improved by post processing (Fig. S2). Increasing the median filter from 5 to 10, 20, and 40 increased valid τ(φ) ROIs from 49.42% to 80.38%, 95.77%, and 99.62%, and increased valid τ(m) ROIs from 66.54% to 92.88%, 98.85%, and 99.81%. Downsampling from the original 512 × 512 pixels to 256 × 256 or 128 × 128 pixels also increased valid ROI recovery to approximately 99.6–99.8%. However, even though median filtering increased the number of valid ROI and reduced τ(φ) variability from an SD of 0.207 at median filter 5 to 0.116 at median filter 40, the variability of τ(m) increased from an SD of 0.226 to 0.417, whereas intermediate- and high-photon acquisition showed lower τ(m) variability (SD 0.125 and 0.130). Downsampling similarly improved valid ROI recovery and reduced τ(φ) variability, but τ(m) remained highly variable. To optimize the single-cell NAD(P)H FLIM workflow, we therefore prioritized high photon-count acquisition (resolution: 512, scan speed: 200, laser power: 5%, line accumulation: 2) for the subsequent image acquisition, as this provided more robust lifetime estimation and helped producing valid cell-specific ROIs.

Bi-exponential fitting returned finite lifetime estimates for all cellular ROIs in the three photon-count groups (Fig. S3; Supplementary Table 9). Mean short-component lifetimes were 0.617–0.633 ns, long-component lifetimes were 3.589–3.752 ns, and the short component contributed approximately 70% of the fitted amplitude. These similar group means indicated that decays pooled over a segmented cell could be fitted across the tested photon-count regimes. However, single-cell τ and τ values were more dispersed in the low-photon group than in the intermediate- and high-photon groups, consistent with the broader phasor distributions at low photon counts. This variability justifies the selected high-photon-count acquisition settings of 512 × 512 pixels, a scan speed of 200 Hz, 5% laser power and 2 line accumulations in application with fitting approach analysis of NAD(P)H lifetime.

We then applied the optimized imaging settings to map the single-cell NAD(P)H intensity and lifetimes along the crypt–villus axis, designating the crypt side as “0” and “1” as the villus (Fig. 1I). We found that NAD(P)H intensity was not equal across different cell types, increasing almost fourfold from 22.35 in the crypt end to a maximum of 85.90 photons/pixel at the villus end. Concomitantly, τ(φ) increased from 1.531 ns in the crypt to 1.715 ns near villus end. τ(m) similarly increased from 2.822 ns to 2.988 ns (Fig. 1J–L), indicating potential differences in reliance on NADH and NADPH-dependent metabolism across the crypt-villus axis.

NAD(P)H-FLIM is frequently offered as an assay performed together with imaging of flavins (FAD, FMN)^12,18^, however, different tissues and cell types may display strong differences in intensity and localization of these pigments^11,34^. We imaged NAD(P)H and flavin-associated autofluorescence in a live porcine small intestinal organoid (Fig. S4A) and measured excitation spectra in two fixed emission bands (Fig. S4B). NAD(P)H emission collected at 420-490 nm peaked at 770 nm excitation, whereas in the 500-600 nm FAD emission channel, the strongest signal occurred with 780 nm excitation, most likely due to the NAD(P)H signal leakage into the 500-600 nm channel. The conventional ‘flavin’ excitation yielded much weaker signals than that of NAD(P)H, being 3.32 at 870 nm and 1.99 a.u. at 900 nm (Fig. S4B). Interestingly, 3D live imaging showed strikingly different intracellular distribution between NAD(P)H (cytoplasm, mitochondria) and FAD (lysosome-like foci), making their combined use of these pigments difficult for our biological model and in the context of metabolic imaging. Collectively, NAD(P)H-FLIM showed high brightness and sufficient cell resolution for subsequent live imaging studies with the small intestinal organoid model, while flavin-like autofluorescence displayed potentially non-relevant for metabolism weak signal.

### Live cell proliferation labeling combined with NAD(P)H-FLIM identify proliferation-associated and differentiated cell-enriched epithelial states

Observation of the drastic increase of cellular NAD(P)H intensity and fluorescence lifetime along the “crypt-villus” axis (Fig. 1I-L) prompted us to investigate whether these gradients could be linked to proliferative and differentiated epithelial states. To test this, we employed a previously developed live proliferation labeling method, based on quenching of Hoechst 33342 fluorescence lifetime by BrdU^35^, together with NAD(P)H-FLIM. Both Hoechst 33342 and NAD(P)H share excitation and emission in multiphoton mode (680∼750 nm) and display different intracellular localization, prompting for a fine-tuning of Hoechst 33342 staining concentration (Fig. 2A). Analysis of the effect of different Hoechst staining concentrations from 50 nM to 2 µM on co-localization and phasor lifetimes (Fig. S5A) revealed 400 nM concentration as optimal for resolving between nuclear and cytoplasmic signals.

**Figure 2.**
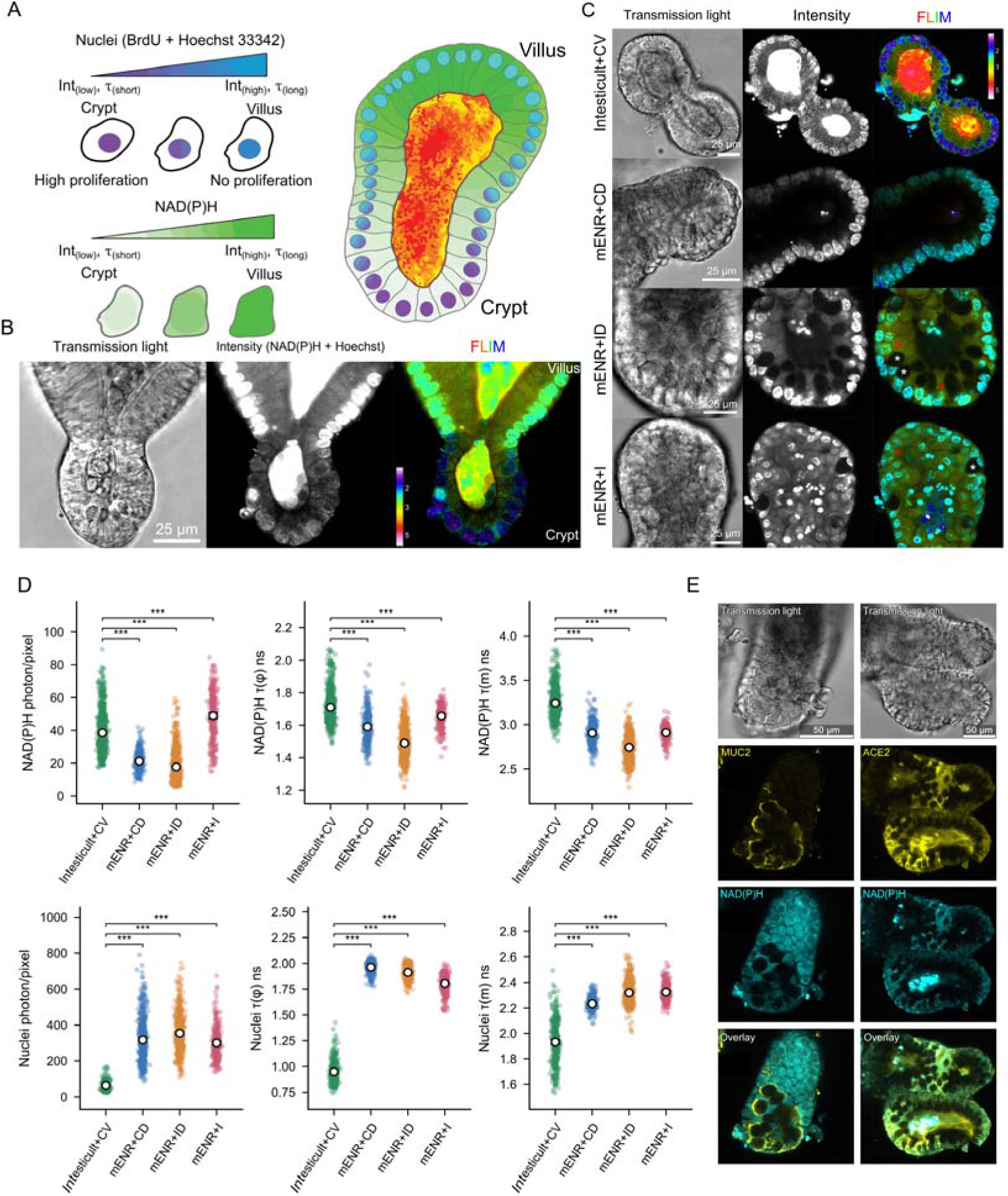
Live cell proliferation labeling, guided differentiation and NAD(P)H-FLIM distinguish between proliferative and differentiated epithelial states in intestinal organoids. **A**: Schematic illustrating nuclear Hoechst 33342 and cellular NAD(P)H signals along the crypt–villus axis. Proliferating cells in the crypt region incorporate BrdU, resulting in Hoechst 33342 quenching with lowering nuclear intensity and shorter fluorescence lifetimes. Non-proliferating differentiated cells in the villus region do not incorporate BrdU, resulting in unquenched Hoechst 33342 signal with high nuclei intensity and long fluorescence lifetime. Cellular NAD(P)H signal changes along the crypt–villus axis, with crypt-like cells showing lower intensity and shorter lifetime, and villus-like cells showing higher intensity and longer lifetime. **B**: Representative transmission light, intensity, and FLIM images showing changes in nuclear Hoechst 33342 and cellular NAD(P)H intensity and lifetime along the crypt–villus axis. FLIM images are displayed as _τ_(m). Scale bar: 25 µm. **C**: Representative transmission light, intensity, and FLIM images of nuclei Hoechst 33342 and cellular NAD(P)H signals upon guided differentiation resulting in distinct intestinal epithelial cell states: stem cell-enriched (IntestiCult + CV), Paneth cell-enriched (mENR + CD), goblet cells (mENR + ID) and enterocyte-enriched (mENR + I). Scale bar: 25 µm. Asterisks indicate high (red) and low (white) NAD(P)H intensity in the cells. **D**: Single-cell quantification of cellular NAD(P)H and nuclei Hoechst 33342 signals across the four guided differentiation conditions. The first row shows cellular NAD(P)H photons/pixel, _τ_(_φ_), and _τ_(m). The second row shows nuclei Hoechst 33342 photons/pixel, _τ_(_φ_), and _τ_(m). **E**: Immunofluorescence validation of organoids differentiated under the mENR + ID condition. ACE2 was used as an enterocyte marker and MUC2 as a goblet cell marker. ACE2-positive regions overlapped with areas of high NAD(P)H signal, whereas MUC2-positive regions overlapped with areas of low NAD(P)H signal. Scale bar: 50 µm. Medium component abbreviations: C: CHIR99021, V: valproic acid, D: DAPT, I: IWP-2. Quantitative summaries and existing comparisons are provided in Supplementary Table 10.

Using this live proliferation labeling method with NAD(P)H-FLIM (Fig. 2A-B), we could now confirm the lower nuclear intensity and lifetimes in proliferative crypt regions, correlating with the metabolic ‘crypt-villus’ gradient (Fig. 1I). This spatially aligned with lower NAD(P)H intensity and shorter lifetime in crypt cells, contrasting cells within the villus region. Thus, we could now relate the proliferation-associated nuclear readout with the cellular NAD(P)H state across cell types in the organoids.

However, the spatial gradient did not identify which epithelial cell states generated the high- and low-NAD(P)H signals. We therefore applied ‘guided differentiation’ organoid culture conditions ^36^ to produce organoids enriched with stem, Paneth, goblet or enterocyte states. First, we required a starting population with a broad diversity of proliferative cells. Under mENR + CV conditions, organoids visualized via BrdU–Hoechst 33342/ NAD(P)H-FLIM retained a subset of unquenched nuclei with high NAD(P)H levels. This pattern indicated the presence of persistent non-proliferating regions in the cultures grown in mENR + CV conditions. We found that using commercial “IntestiCult” media supplemented with CV displayed stronger and more uniformly distributed cell proliferation, evidenced by Hoechst 33342 quenching (Fig. S5B). Furthermore, with the culture of Lgr5-GFP organoids^2^, IntestiCult + CV medium also resulted in stronger and more broadly distributed GFP fluorescence than mENR + CV medium. Under mENR + CV, high Lgr5-GFP expression remained confined mainly to crypt-like regions (Fig. S5C). IntestiCult + CV was therefore selected as the stem cell-enriching starting growth condition.

After 2–3 days of growth in IntestiCult + CV, organoids were transferred to three mENR-based differentiation media. mENR + CD, mENR + ID and mENR + I were used for Paneth, goblet cell and enterocyte enrichment, respectively (Fig. 2C).

Two imaging workflows were used to quantify cellular NAD(P)H and nuclear signals: BrdU–Hoechst 33342/ NAD(P)H-FLIM was used to characterize nuclear intensity and lifetime (Fig. S6B). However, Hoechst 33342-associated population remained on the phasor plot even after the nuclear regions were cropped from the images. This residual signal prevented reliable isolation of the cellular NAD(P)H from the BrdU–Hoechst dataset. We therefore used a separate NR12S/ NAD(P)H dataset for cellular segmentation and cellular NAD(P)H comparison (Fig. S6A). The four culture conditions produced distinct single-cell NAD(P)H distributions. Mean cellular intensities were 38.54 photons/pixel in IntestiCult + CV (Stem cell); 21.27 in mENR + CD (Paneth cell); 17.69 in mENR + ID (Goblet cell) and 48.87 in mENR + I (Enterocytes). Concomitantly, mean τ(φ) were 1.710, 1.591, 1.489 and 1.656 ns; mean τ(m) were 3.243, 2.905, 2.743 and 2.910 ns. Among the three mENR-based conditions, mENR + I medium (inducing enterocytes differentiation) had the highest mean NAD(P)H intensity and both lifetime measures. Meanwhile, nuclear Hoechst intensity was quenched with 63.05 photons/pixel in IntestiCult + CV and increased to 316.72, 353.91 and 299.94 photons/pixel after CD, ID and I differentiation. Nuclear Hoechst τ(φ) and τ(m) increased from 0.949 ns to 1.962, 1.911, 1.805 ns and 1.932 to 2.232, 2.319, 2.324 respectively after CD, ID and I differentiation (Fig. 2D).

However, the longer NAD(P)H lifetime in IntestiCult + CV conditions appeared inconsistent with the shorter lifetime observed in our earlier single-cell mapping (Fig. 1I). Since we used IntestiCult for stem-cell induction and mENR for subsequent lineage differentiation, differences in medium composition could contribute to this discrepancy. We therefore compared NAD(P)H fluorescence across culture and imaging-medium conditions (Fig. S6C, D). Organoids cultured in IntestiCult or mENR were imaged directly in the same medium, whereas the imaging-medium (IM; phenol-red-free DMEM supplemented with D-glucose, GlutaMAX, sodium pyruvate and HEPES-Na; see *Methods*) group was transferred from mENR shortly before imaging. NAD(P)H τ(φ) was 1.957 ns in IntestiCult, 1.667 ns in mENR and 1.709 ns in imaging medium (n = 7, 5 and 8 organoids). Corresponding τ(m) values were 3.509, 3.034 and 3.058 ns. The longer lifetimes in the IntestiCult group were consistent with a medium-associated contribution to the elevated NAD(P)H lifetimes in IntestiCult + CV, although this comparison did not separate culture history from the medium present during imaging.

Double-exponential fitting of fluorescence lifetimes reproduced the broad range of observed phasor lifetimes while resolving contributions from component lifetimes and amplitudes (Fig. S7; Supplementary Table 11): IntestiCult + CV conditions had the longest mean long-component lifetime, whereas mENR + ID had the shortest and the largest short-component amplitude fraction. The amplitude-weighted lifetime followed the same group ordering as τ(φ): IntestiCult + CV (stem-cell enriched) > mENR + I (enterocyte enriched) > mENR + CD (Paneth-cell enriched) > mENR + ID (goblet-cell enriched). This ordering was consistent with the longer NAD(P)H lifetimes associated with IntestiCult culture. mENR + ID and mENR + I medium grown organoids displayed presence of distinct high- and low-NAD(P)H cell populations (Fig. 2C, indicated with asterisks). High-NAD(P)H cells were more prominent after mENR + I treatment, consistent with enrichment of enterocytes in the culture. To determine their cellular identity, we combined NAD(P)H imaging with immunofluorescence for ACE2 and MUC2 under mENR+ID growth condition. ACE2-positive enterocytes colocalized with high-NAD(P)H regions, whereas MUC2-positive goblet cells colocalized with low-NAD(P)H regions (Fig. 2E). These marker-defined colocalization patterns confirmed that the enterocytes represented the ‘high-NAD(P)H’ cells, whereas goblet cells exhibited lower NAD(P)H fluorescence. We further examined this association in organoids carrying the Muc2-iCre × Rosa26-YFP reporter^37^. We found that during enterocyte differentiation, high-NAD(P)H regions did not overlap with Muc2-YFP-positive cells: instead, Muc2-YFP-positive goblet cells were associated predominantly with low NAD(P)H signals. Additionally, Muc2-YFP fluorescence was markedly reduced following Paneth cell differentiation but remained abundant upon goblet cell directed differentiation (Fig. S8A). Together, the immunofluorescence and use of fluorescent reporter organoid line identified enterocytes as the principal high-NAD(P)H cell population in mouse small intestinal organoids.

To achieve deeper insight in changes of NAD(P)H metabolism during Paneth and goblet cell development, we performed staining with Zinpyr-1 dye^38^, labeling Paneth cells (Fig. S8B). Consistent with previous reports, Zinpyr-1 showed bright fluorescence in crypt regions of normal mouse small intestinal organoids and in Paneth cell-enriched organoids. However, it was also detected in some cells under the goblet cell-enriched condition (Fig. S8B). Zinpyr-1 therefore preferentially highlighted strongly Zn-positive cells but did not identify Paneth cells exclusively. Both differentiation conditions showed BrdU-quenched nuclei and bright NAD(P)H regions on day 1, followed by increasing numbers of unquenched nuclei through days 2–3 (Fig. S8C, D).

### Efflux inhibition and mitochondrial imaging indicate higher mitochondrial membrane potential in crypt than in villus regions

Having identified enterocytes as the high-intensity, long-lifetime NAD(P)H population along the crypt-villus axis, we wondered if these changes could be correlated to mitochondrial membrane potential^39–41^. Using labeling of active cell mitochondria with the tetramethylrhodamine (TMRM), we unexpectedly found its fluorescence intensity enriched in crypts and weak in villus regions (i.e. potentially indicating decrease of ΔΨm), contrasting the increase of NAD(P)H fluorescence towards the villus (Fig. 3A). Across 50 normalized spatial bins, TMRM decreased from 110.4 to 34.9 photons/pixel but NAD(P)H increased from

**Figure 3.**
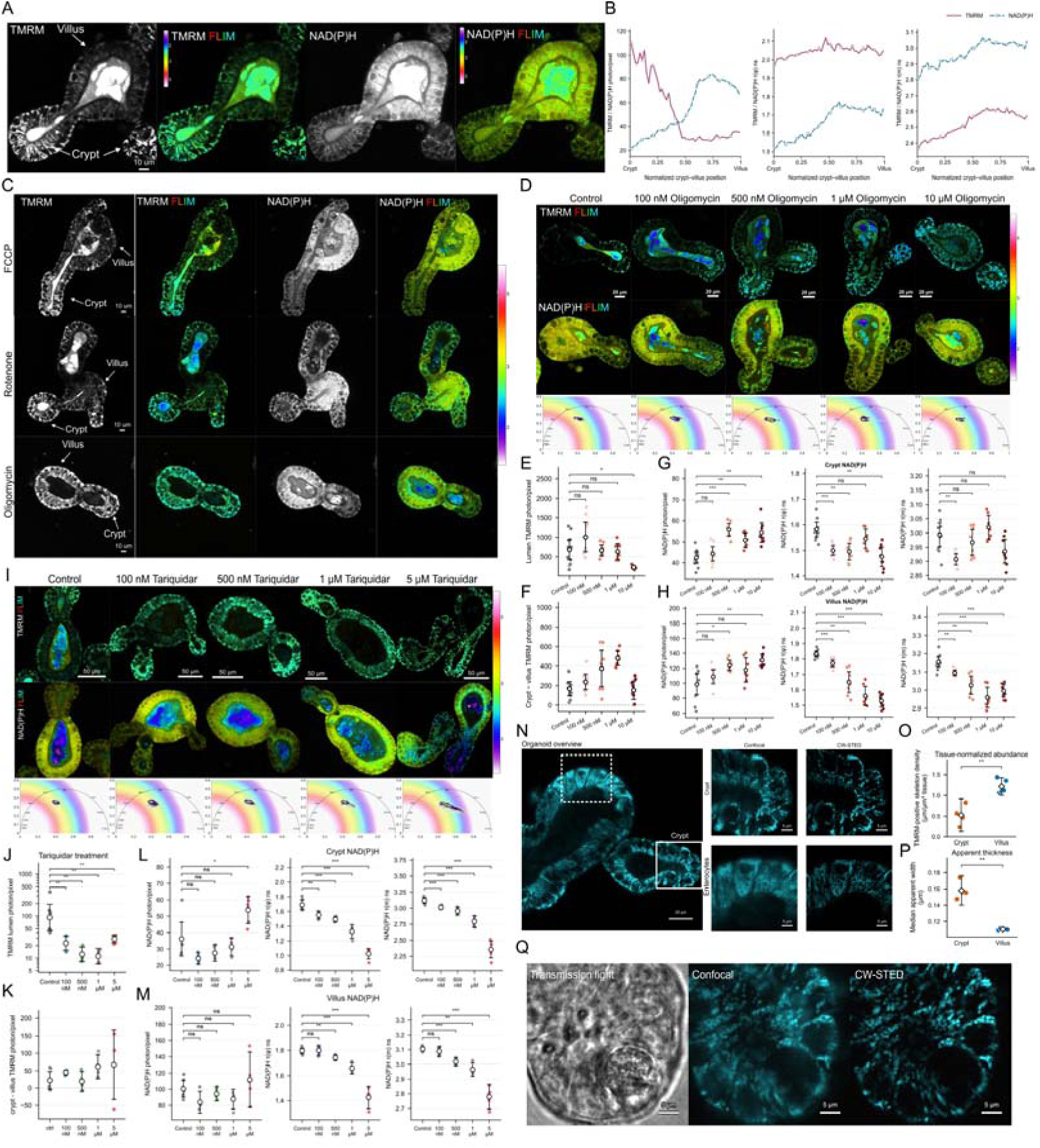
Comparison between regional mitochondrial polarization, organization and NAD(P)H distribution in live mouse intestinal organoids. **A**: Representative TMRM and NAD(P)H intensity and fluorescence lifetime imaging microscopy (FLIM) images of live mouse intestinal organoids stained with 10 nM TMRM. Arrows indicate crypt and villus regions, displaying crypt-enriched TMRM fluorescence and villus-enriched NAD(P)H fluorescence. **B**: Spatial profiles of TMRM and NAD(P)H intensity, _τ_(_φ_), _τ_(m), along the normalized crypt–villus axis, with crypt and villus positions designated 0 and 1, respectively. Profiles show descriptive pooled pixel means across 50 spatial bins. **C**: Representative intensity and FLIM images following 2 µM FCCP, 10 µM rotenone or 10 µM oligomycin treatment. **D**: Representative TMRM and NAD(P)H-FLIM images and corresponding NAD(P)H phasor plots under control and oligomycin treatment conditions at 100 nM, 500 nM, 1 _μ_M or 10 _μ_M. **E–H**: Quantification of luminal TMRM intensity **(E)**, paired crypt-minus-villus TMRM intensity differences **(F)**, and NAD(P)H intensity, _τ_(_φ_) and _τ_(m) in crypt **(G)** and villus **(H)** regions; n = 11, 9, 8, 7 and 8 images for control and increasing oligomycin concentrations, respectively. **I**, Representative TMRM and NAD(P)H FLIM images and corresponding NAD(P)H phasor plots following tariquidar treatment at 100 nM, 500 nM, 1 _μ_M or 5 _μ_M. **J–M**: Quantification of luminal TMRM intensity **(J)**, paired crypt-minus-villus TMRM intensity differences **(K)**, and NAD(P)H intensity, _τ_(_φ_) and _τ_(m) in crypt **(L)** and villus **(M)** regions. n = 7, 5, 5, 5 and 6 images for control and increasing tariquidar concentrations; Positive differences in F,K indicate greater crypt TMRM accumulation. **N**: TMRM-labelled organoid overview and corresponding confocal and continuous-wave stimulated emission depletion (CW-STED) images of crypt and villus-enterocyte regions; boxes identify the enlarged regions. **O**, Tissue-normalized TMRM-positive mitochondrial skeleton density, calculated as total skeleton length divided by tissue area. **P**, Median apparent mitochondrial width derived from segmented CW-STED images. For **O,P**, n = 4 crypt and 5 villus fields; **Q**: Transmission light, confocal and CW-STED images of a crypt containing a Paneth cell, identified by prominent secretory granules, and surrounding putative stem cells. Quantitative summaries and existing comparisons are provided in Supplementary Table 12.

21.4 to 66.9 photons/pixel. NAD(P)H τ(φ) and τ(m) increased from 1.52 to 1.73 ns and 2.81 to 3.04 ns, respectively; TMRM lifetimes also modestly increased, indicating decrease of mitochondrial polarization towards the villus tip (Fig. 3B). To understand the discrepancy between NAD(P)H fluorescence and TMRM mitochondrial accumulation, we analyzed publicly available single-cell RNA dataset for mouse intestinal organoids^42,43^. At the level of transcriptome, enterocytes showed the highest mean values for enzymes involved in TCA-cycle, Electron Transport Chain (ETC) complexes I–IV and ATP-synthase expression scores among the four compared lineages. More specifically, ATP synthase (Complex V) gene sets were also highly expressed in enterocytes (Supplementary Fig. S9A-C). These signatures motivated us to further investigate whether ATP-synthase-mediated proton utilization could be the cause of the lower steady-state membrane potential.

We therefore interrogated organoids with drugs affecting mitochondrial function: FCCP (mitochondrial uncoupler: 2 μM), rotenone (complex I inhibitor: 10 μM) or oligomycin (complex V inhibitor: 10 μM). Interestingly, oligomycin visibly increased villus TMRM fluorescence (Fig. 3C), although the initial villus region-level comparison showed a nonsignificant intensity increase. Meanwhile, villus TMRM τ(φ) and τ(m) decreased, whereas crypt decreases were nonsignificant under oligomycin treatment. Rotenone and oligomycin shortened crypt and villus NAD(P)H lifetimes; while FCCP produced no differences in this screen (Fig. S9 D–G). Together, villus TMRM signal recovery after ATP synthase blocking suggested that enterocytes could utilize proton gradient more efficiently than stem cells. However, luminal TMRM signal complicated such bioenergetic explanation: 10 μM oligomycin also markedly reduced luminal TMRM fluorescence, although TMRM remained in the culture medium. TMRM accumulation depends on mitochondrial and plasma membrane potentials and can also be influenced by multidrug transporter activity^44,45^. The fall in luminal fluorescence therefore prompted us to examine apical dye efflux. Previous reports on oligomycin inhibition of P-glycoprotein-mediated transport provided a rationale for investigating this possibility^46–48^. We therefore examined the dose-dependent effect of oligomycin on the TMRM dye accumulation in the lumen (Fig. 3D). Cellular responses to oligomycin were evident at lower concentrations: NAD(P)H lifetimes decreased in both crypt and villus regions from 100 nM (Fig. 3G,H), while TMRM intensity generally increased and its lifetimes shortened across the concentration series (Supplementary Fig. S9H, I). Villus TMRM intensity was significantly elevated at 1 μM, increasing approximately threefold. By contrast, luminal TMRM decreased significantly only at 10 μM, falling by 67% relative to control (Fig. 3E). Thus, detectable cellular TMRM responses resulted in luminal signal depletion, suggesting that high dose of oligomycin could influence ATP-dependent apical efflux^48^. Nevertheless, crypt TMRM intensity remained higher than villus intensity throughout the series (Fig. 3F). We therefore tested apical efflux inhibitors to determine whether apical efflux contributed to the low villus TMRM signal and whether crypt-enriched accumulation persisted after efflux suppression, helping to clarify its relationship with observed regional (crypt/ villi) mitochondrial membrane potential differences.

Before reassessing TMRM distribution, we first tested the effect of efflux inhibition on luminal accumulation of rhodamine 123, an established transepithelial transport probe. Its apical transport into intestinal-organoid lumen is sensitive to P-glycoprotein inhibition^49–51^. Treatment with tariquidar (5 μM) and oligomycin (10 μM) reduced mean luminal fluorescence from 113.3 to 19.0 and 15.4 photons/pixel, respectively (Fig. S9 J,K). We also found that crypt-adjacent luminal region showed significant longer τ(φ) and τ(m) values than villus-adjacent with no drastic change of the intensity (Fig. S9 L–N). Notably, Rhodamine 123 did not reproduce the TMRM crypt-high/villus-low mitochondrial pattern since its working concentration (5 μM) normally exceeded the reported non-quenching range of 10–40 nM^41^, limiting direct interpretation of its intensity as steady-state mitochondrial membrane potential; at the same time TMRM at 10 nM fell within this range and was therefore better suited to regional mitochondrial membrane potential comparisons^52,53^. After validating tariquidar as an effective apical efflux inhibitor in organoids, we next performed TMRM–NAD(P)H imaging following tariquidar treatment at 0.1–5 μM. We observed a clear recovery of villus TMRM labeling from 100 nM, accompanied by markedly reduced luminal fluorescence (Fig. 3I,J). Nevertheless, mean crypt TMRM intensity remained higher than in the villus across the range of concentrations (Fig. 3K). Thus, we found that the apical efflux process contributed to the observed low baseline enterocyte TMRM signal, but its inhibition did not eliminate crypt-confined TMRM accumulation. Such residual difference could reflect either difference in mitochondrial polarization or in mitochondrial number. Increasing tariquidar concentrations also resulted in shorter NAD(P)H τ(φ) and τ(m) in both crypt and villus regions (Fig. 3L, M), potentially indicating high metabolic demand required for the efflux transporters. To determine whether NAD(P)H lifetime shortening reflected apical efflux inhibition rather than a tariquidar-specific side effect, we tested another efflux inhibitor, elacridar. Luminal TMRM loss became evident from 500 nM (Fig. S10A,B). Crypt TMRM intensity nevertheless remained higher than villus intensity across the concentration series (Fig. S10C). NAD(P)H lifetimes also shortened with increasing elacridar concentration, particularly in villus regions (Fig. S10D,E). Thus, the use of two apical efflux inhibitors supported a link between efflux inhibition and effect on NAD(P)H lifetime. Notably, at 5 μM, both inhibitors produced bright, short-lifetime puncta in the NAD(P)H detection channel within crypt regions (Fig. S10F). These features were more prominent with tariquidar and could contribute to the increased intensity and variability observed at this concentration.

We also performed time-lapse TMRM–NAD(P)H imaging after adding 500 nM tariquidar or 10 μM oligomycin following baseline acquisition. Both treatments increased villus TMRM fluorescence while reducing luminal labeling (Fig. S10G,J and Supplementary Videos 1 and 2). After normalization to each image series baseline (F/F), villus TMRM reached 3.05-fold baseline over 20.8 min with tariquidar and 2.09-fold over 18.6 min with oligomycin (Fig. S10H,K). Despite this shared TMRM redistribution, the accompanying NAD(P)H responses differed (Fig. S10I,L). Tariquidar produced minor change in NAD(P)H intensity, which reached 1.02-fold compared to baseline at the final frame. Oligomycin increased NAD(P)H intensity to a peak of 1.14-fold at 8.3 min, followed by a partial decline to 1.09-fold. Oligomycin and tariquidar influenced NAD(P)H lifetime similarly, leading to the shortening of both τ(φ) and τ(m): NAD(P)H τ(φ) decreased from 1.71 to 1.67 ns with tariquidar and from 1.77 to 1.54 ns upon oligomycin addition. This data points at difference between oligomycin and tariquidar effects on organoids metabolism, which might reflect their efflux inhibitory action.

To determine whether persistent crypt TMRM enrichment after efflux inhibition reflected in greater mitochondrial polarization or a denser mitochondrial network, we visualized TMRM-labelled mitochondrial organization using confocal and CW-STED imaging (Fig. 3N). Crypt-based mitochondria appeared less densely distributed and thicker than those present in villus enterocytes. For quantification, mitochondrial masks generated with Nellie^54^ were skeletonized, and total skeleton length was normalized to tissue area, excluding luminal spaces (Supplementary Fig. S10M,N). Villus regions showed 2.32-fold higher TMRM-positive skeleton density than crypts (Fig. 3O). Conversely, mean field-level median apparent mitochondrial width was greater in crypts than in villi (0.158 versus 0.110 μm; Fig. 3P), with local width estimated as twice the skeleton-to-mask-background distance (Supplementary Fig. S10O). To identify the cellular source of the strong crypt TMRM signal, we compared a Paneth cell with neighboring stem cells. The Paneth cells, identified by prominent secretory granules in transmission light, showed weaker mitochondrial labelling. Consistent with previous reports^22^, this suggests that stem cells and highly proliferative daughter cells are the main contributors to the strong crypt TMRM signal. (Fig. 3Q). Together, persistent crypt TMRM high signal upon efflux inhibition and the lower measured crypt mitochondrial density support the higher mitochondrial membrane potential in crypt stem cells than in villus enterocytes. Enterocytes instead contained denser and thinner TMRM-positive mitochondrial network, potentially contributing to their higher NAD(P)H intensity and longer lifetimes.

### Villus regions demonstrate high mitochondrial ATP, whereas crypts show high levels of ROS

Our finding of higher ΔΨm / short NAD(P)H lifetimes in crypts, contrasting villus regions, prompted us to investigate how such differences could reflect mitochondrial production of ATP. However, conventional ATP assays offer only a bulk readout, providing an average across metabolically distinct, dynamic cell populations and obscuring regional differences^6,55^. Although guided organoid differentiation can help enriching the selected lineages, it also changes the original cellular composition, interfering with the spatial organization of the crypt–villus epithelium and potentially the ‘metabolic phenotype’ (Fig. 2, S6). We therefore tested ATP-Red 1 dye as a probe to visualize cell-specific ATP dynamics in cells subjected to mitochondrial inhibition within the living organoids.

ATP-Red 1 was originally reported as an ATP-responsive, predominantly mitochondria-localizing fluorescence intensity probe ^55–57^. With the small intestinal organoids culture, we found that ATP-Red 1 provided bright staining with higher intensity in crypts than in villus regions, in addition to pronounced luminal fluorescence (Fig. 4A), supporting the idea that ABC-transporter mediated efflux of ATP-Red 1 (a derivative of rhodamine) can affect the fluorescence intensity read-out, similarly to observed effects of efflux on TMRM (Fig. 3). However, this staining pattern also mirrored our finding on mitochondrial polarization gradient across crypt-villus axis (Fig. 3), revealing membrane potential-dependent accumulation and apical efflux effects for ATP-Red 1. We therefore tested whether the fluorescence lifetimes of ATP-Red 1 could help visualize the ATP pool in organoids. We found that lower fluorescence intensity villus regions exhibited longer ATP-Red 1 lifetimes, producing distinct regional distributions, compared to the crypt, on the phasor plot (Fig. 4B). Across seven paired images, mean intensity was higher in crypts than in villus regions (46.3 versus 30.2 photons/pixel). Conversely, villus ATP-Red 1 τ(φ) was 0.203 ns longer (2.305 versus 2.102 ns). τ(m) was also 0.410 ns longer (2.965 versus 2.555 ns; Fig. 4C).

**Figure 4.**
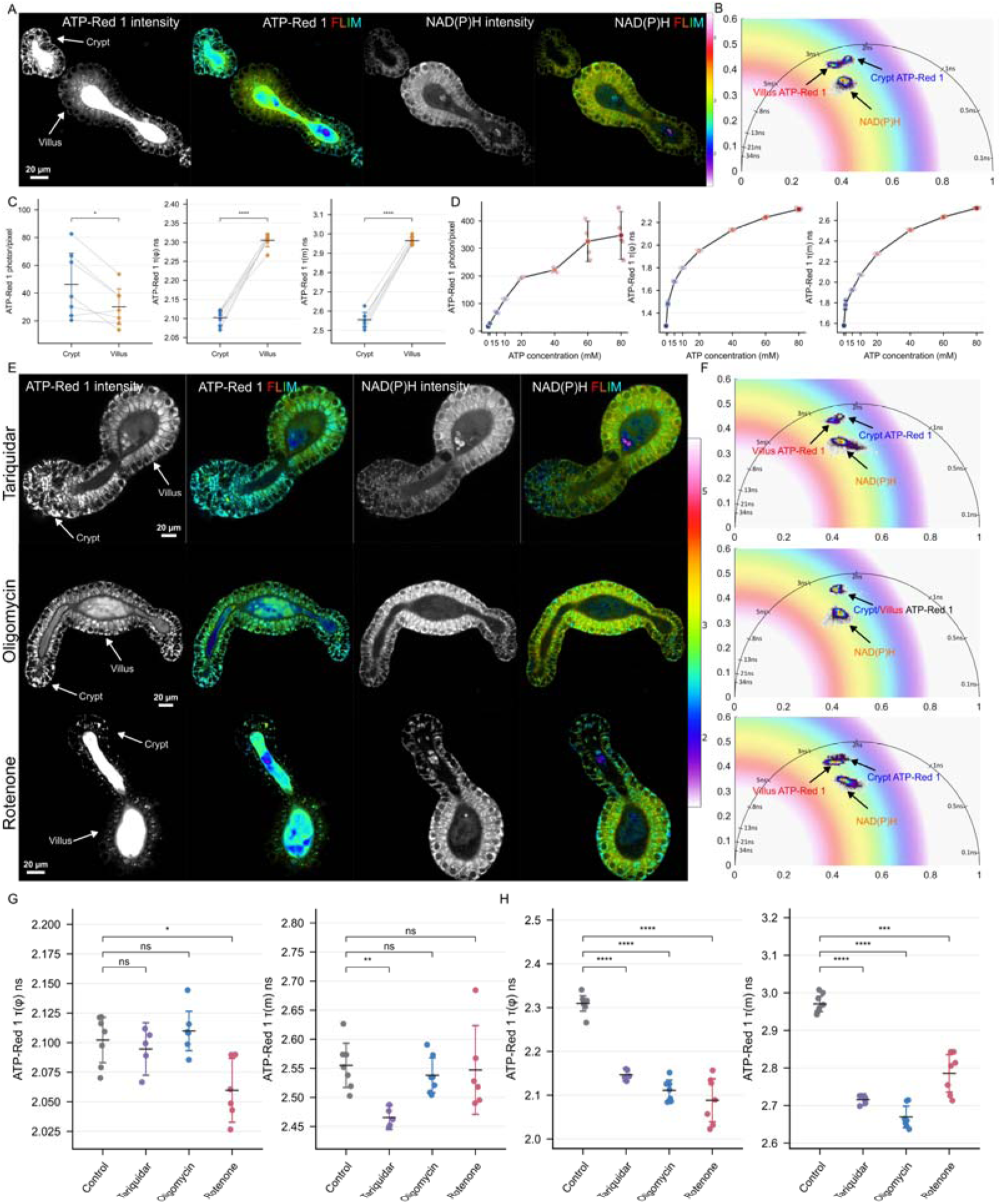
Regional ATP-Red 1 fluorescence and lifetime responses to mitochondrial and efflux inhibition in live mouse intestinal organoids. **A**: Representative ATP-Red 1 and NAD(P)H intensity and FLIM images under control conditions. Arrows indicate crypt and villus regions. ATP-Red 1 fluorescence is enriched in crypts and lumen, whereas NAD(P)H fluorescence is higher in villus regions. **B**: Corresponding phasor plot showing crypt and villus ATP-Red 1 distributions and NAD(P)H fluorescence. **C**: Paired comparisons of ATP-Red 1 intensity, _τ_(_φ_), and _τ_(m); n = 7. Lines connect regions from the same acquisition. **D**: ATP-Red 1 intensity and lifetimes measured at 0, 1, 5, 10, 20, 40, 60 and 80 mM ATP in 25 mM HEPES buffer, pH 7.8; n = 5 acquisition images per concentration. Connecting lines show the concentration-dependent mean response. **E**: Representative ATP-Red 1 and NAD(P)H intensity and FLIM images following tariquidar (500 nM), oligomycin (10 _μ_M) or rotenone (10 _μ_M) treatment. Arrows indicate crypt and villus regions. **F**, Corresponding phasor plots, ordered as in E. **G,H**: ATP-Red 1 phase and modulation lifetimes in crypt **(G)** and villus **(H)** regions. Quantitative summaries and existing comparisons are provided in Supplementary Table 13.

To better understand the relationship between ATP concentration and ATP-Red 1 lifetime we calibrated ATP-Red 1 across 0–80 mM ATP (Fig. S11A). Mean intensity increased from 17.2 to 347.9 photons/pixel across this range. τ(φ) and τ(m) increased from 1.287 to 2.317 ns and from 1.580 to 2.719 ns, respectively (Fig. 4D). The corresponding phasor trajectory shifted towards longer lifetimes (Fig. S11B). These measurements supported an ATP-responsive lifetime readout for ATP-Red 1, correlating the longer villus lifetime with higher ATP level. We next validated the ATP sensitivity of this dye in FLIM, using treatment of organoids with tariquidar (500 nM), oligomycin (10 μM) and rotenone (10 μM), targeting apical efflux, ATP synthase and respiratory complex I, respectively. Luminal ATP-Red 1 fluorescence disappeared after tariquidar and oligomycin treatment but remained prominent after rotenone treatment (Fig. 4E). Oligomycin-induced loss of luminal ATP-Red 1 fluorescence was highly similar to what we observed with TMRM (Fig. 3). Both tariquidar and oligomycin treatments led to an increase in villus-specific ATP-Red 1 intensity. However, if we interpret this increase solely as an ATP-linked response, it would imply greater ATP availability while ATP synthase is inhibited. Collectively, these observations confirmed poor suitability of the ATP-Red 1 dye-based intensity imaging of ATP. Since we confirmed the reliability of a lifetime readout of ATP-Red 1, the following analysis was focused on comparison of the fluorescence lifetime parameter of this probe.

Treatment of organoids with oligomycin led to the concomitant response in ATP-Red 1 fluorescence lifetime (Fig. 4F–H): villus ATP lifetime τ(φ) decreased from 2.310 to 2.111 ns, and τ(m) from 2.970 to 2.670 ns, whereas crypt lifetimes remained essentially unchanged (2.102 versus 2.110 ns for τ(φ) and 2.555 versus 2.538 ns for τ(m), both non-significant). The greater lifetime shortening (reflecting decrease in ATP) in villus regions suggested that ATP pool in these regions was more dependent on mitochondrial ATP synthase activity. Rotenone-treated villus regions also decreased τ(φ) and τ(m) up to 2.088 and 2.786 ns correspondingly (Fig. 4H). In crypts, τ(φ) decreased modestly from 2.102 to 2.060 ns, whereas τ(m) remained unchanged (Fig. 4G). Tariquidar-treated villus regions likewise showed shorter lifetimes than the control (τ(φ): 2.147 ns; τ(m): 2.715 ns, Fig. 4H). However, the crypt ATP response was lifetime-metric-dependent: τ(φ) remained unchanged at 2.095 ns, whereas τ(m) decreased to 2.465 ns (Fig. 4G). The villus ATP-Red 1 lifetime shortening accompanied the NAD(P)H lifetime shortening previously observed under tariquidar treatment, raising the possibility that apical efflux inhibition also influenced mitochondrial metabolism and ATP availability. A link between mitochondrial ATP supply and ABC transporter activity has also been demonstrated in chemoresistant cancer cells, where mitochondrial ATP supported drug efflux^48^. Furthermore, ATP-Red 1 rotenone and oligomycin responses were supported by higher electron transport chain (ETC) and ATP-synthase-related gene expression, as well as denser mitochondrial networks that we found in enterocytes (Fig. 3N–P, S9A–C). Collectively, this data suggests that villus regions depend on oxidative phosphorylation more strongly than crypts. This interpretation aligns with previous work of linking intestinal differentiation to increased mitochondrial pyruvate metabolism^58^ and active TCA cycle in enterocytes^24^, which could, in turn, explain the high NAD(P)H fluorescence observed in villus enterocytes. More importantly, the similarly strong effect of oligomycin / tariquidar on mitochondrial ATP production suggests that ATP-dependent function of ABC-transporters (unlike in the crypt region, where stimulations did not show systematic correlative changes in τ(φ) and τ(m) of ATP-Red 1, Fig. 4G–H) can represent one of the main consumers for mitochondrially produced ATP in enterocytes.

The lower ATP detected in crypts prompted us to question the functional role of their higher mitochondrial polarization. Previous research revealed that mitochondrial ROS–p38 signaling contributes to intestinal stem cell differentiation^22^. This raised the possibility that highly polarized mitochondria in crypt-based stem cells could support ROS production and signaling involved in differentiation. We therefore performed ROS labeling in live organoids, using CellROX Deep Red dye and found the higher (almost 8fold) staining in the crypts than in villus regions (79.4 versus 11.7 photons/pixel). Interestingly, CellROX dye fluorescence lifetime also differed, potentially reflecting its chemical transformation across different cells and ROS species (Fig. S12 A,B). We also found that crypt-enriched CellROX staining persisted during co-imaging together with TMRM and NAD(P)H (well-resolved via phasor FLIM) upon efflux inhibition conditions (Fig. S12C–H). Upon treatment with apical efflux inhibitors, the difference of ROS staining with CellROX was ∼threefold between crypt and villus regions. Notably, this apparent reduction in crypt CellROX fluorescence was observed during co-staining with TMRM. A potential explanation for this is in membrane polarization-dependent accumulation of CellROX. Indeed, Kuběnová and colleagues reported loss of mitochondria-associated CellROX Deep Red fluorescence after valinomycin treatment in *Arabidopsis* root hairs^59^, providing evidence for this explanation. Alternatively, direct competition between CellROX and TMRM dyes upon staining can provide additional explanation for this phenomenon. A minor uncoupling effect associated with mitochondrial TMRM accumulation could also reduce the oxidation-sensitive signal.^60^ Despite such intensity responses and potential artifacts, crypt and villus populations remained distinguishable in the CellROX and NAD(P)H phasor FLIM plots, and CellROX lifetimes remained longer in villus regions. To confirm the increased ROS production in crypts, we also labeled organoids with ROSstar 550 (hydro-Cy3) dye, previously used with different ex vivo and in vitro models, including the colon^61,62^: with this dye, we also observed crypt-enriched ROS staining under control (79.3 versus 32.4 photons/pixel) and tariquidar treatment (164.8 versus 49.7 photons/pixel; Fig. S12I–L) conditions. Altogether, our data indicates that increased mitochondrial polarization in crypt-based stem cells does not result in increased ATP production. Instead, these “hyper-polarized” mitochondria are involved in ROS production, potentially representing one of their essential physiological functions, in agreement with previous findings^22,63,64^. Importantly, clear accumulation of ROS in crypts was also reported in mouse intestinal tissue cryosections using an alternative dihydroethidium (DHE) oxidation assay^65^, supporting our results obtained in the live organoid model. Lower fluorescence in villus regions could also reflect more effective antioxidant clearance, even if ROS production remained substantial. In this context, the high NAD(P)H intensity (i.e. its reduced form) in villus region raises the possibility that NADPH-dependent antioxidant systems contributed to ROS removal in enterocytes. Our findings also indicate a regional difference in the net ROS oxidation-sensitive signals.

### Intestinal epithelial cell populations exhibit distinct mitochondrial NAD(P)H responses to respiratory inhibition treatments

Our previous measurements revealed villus-enriched NAD(P)H fluorescence and higher cellular ATP, contrasting with higher mitochondrial membrane potential and ROS levels in the crypt region (Fig. 3, 4). However, these results only indicated ‘steady-state’ equilibrium conditions, which provide only limited information about the ‘flux’ and dynamics of these phenomena. This prompted us to examine the dynamics of cellular responses in NAD(P)H-FLIM, occurring upon application of drugs affecting the electron transport chain and ATP utilization. We measured NAD(P)H intensity, τ(φ) and τ(m), while applying segmentation of polarized mitochondria, using TMRM-staining.

Our previous differentiation experiments revealed differences in Lgr5–GFP-positive cell numbers and NAD(P)H signals between mENR and IntestiCult-grown organoid cultures supplemented with CHIR99021 and valproic acid (CV): IntestiCult + CV-grown organoids showed stronger Lgr5–GFP fluorescence (Fig. S5C) and higher proliferation, as indicated by BrdU-mediated Hoechst quenching (Fig. S5B). We therefore examined whether such induction conditions affected stem cell mitochondrial NAD(P)H pool responses to rotenone and oligomycin. To reduce potential medium-related effects on NAD(P)H lifetime (Fig. S6C, D), the IntestiCult induction protocol comprised 2 days in IntestiCult + CV followed by 2 days in mENR, and then compared with mENR + CV-grown cultures (Fig. 5A–D). For quantification, we combined intermediate- and high intensity-GFP regions defined by per-image intensity percentiles and overlayered these regions with TMRM-derived mitochondrial segmentation masks. This selection retained approximately 75% of common valid GFP pixels and provided an operational definition for the GFP-enriched stem-cell regions.

**Figure 5.**
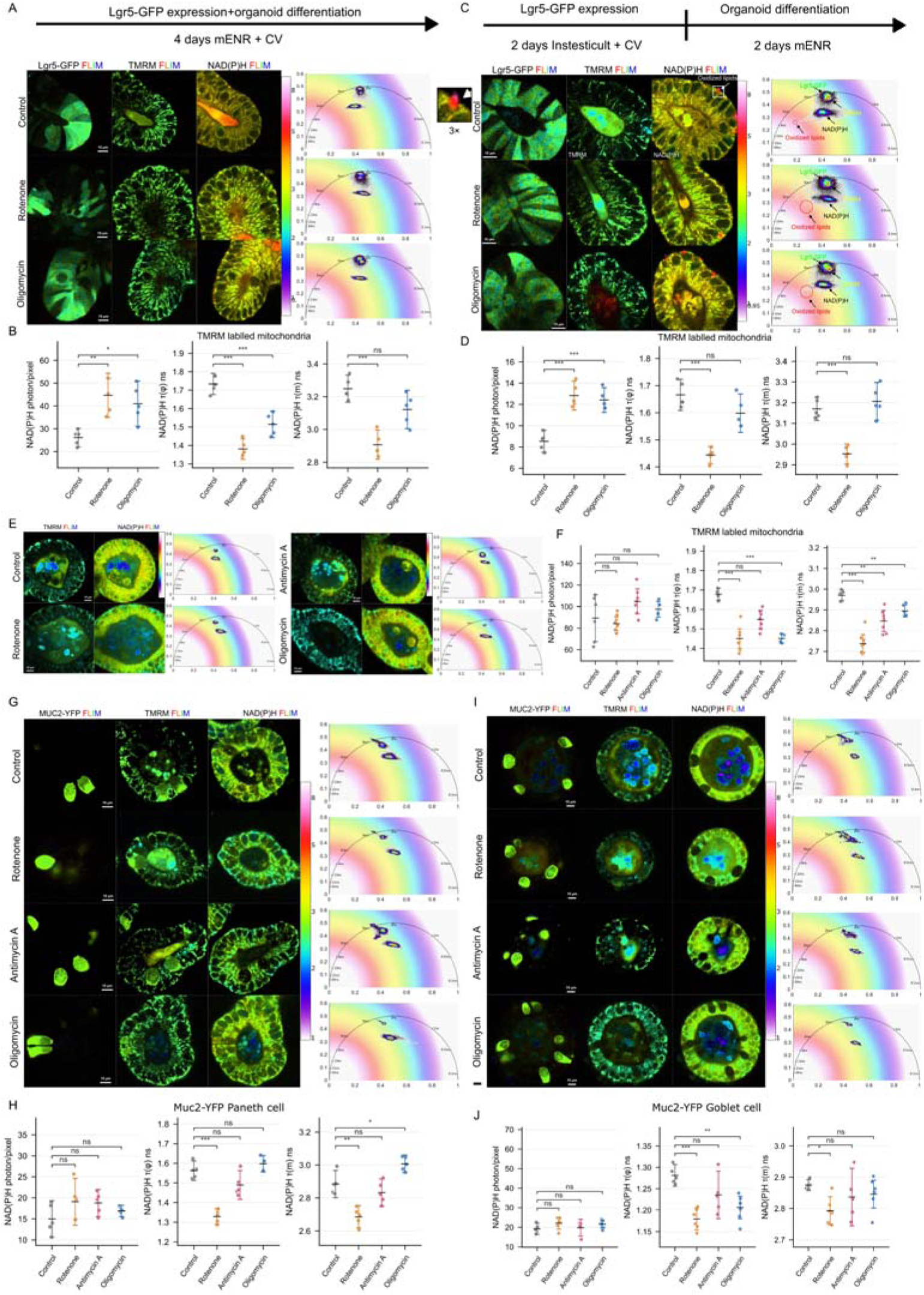
**Cell-type-dependent NAD(P)H responses to inhibition of electron transport and ATP synthesis. A,B**: Representative FLIM images of Lgr5-GFP, TMRM and NAD(P)H with corresponding phasor plots **(A)**, and NAD(P)H intensity, _τ_(_φ_), and _τ_(m) **(B)**, in organoids cultured for 4 days in mENR supplemented with CHIR99021 and valproic acid (CV), under control conditions or following rotenone or oligomycin treatment. NAD(P)H data were extracted from TMRM-positive mitochondrial masks intersected with GFP-enriched stem-cell regions. Intermediate- and high-GFP regions were combined using per-image intensity percentiles, with p25 defining the low-GFP boundary and p75 defining high-GFP seeds, retaining approximately 75% of common valid pixels. **C,D**: Corresponding FLIM images and phasor plots **(C)** and NAD(P)H measurements using the same regional selection **(D)** in organoids cultured for 2 days in IntestiCult + CV followed by 2 days in mENR. In C, the white box and 3× inset highlight a red–orange cytoplasmic punctum in the control NAD(P)H image. These puncta and the additional long-lifetime phasor population are consistent with oxidized-lipid-associated autofluorescence. **E,F**: Representative TMRM and NAD(P)H FLIM images and corresponding phasor plots **(E)**, and NAD(P)H intensity, _τ_(_φ_) and _τ_(m) within TMRM-positive mitochondrial regions of villus enterocytes **(F)**, under control conditions or following rotenone, antimycin A or oligomycin treatment. **G,H**: Representative Muc2-YFP, TMRM and NAD(P)H-FLIM images and corresponding phasor plots of crypt regions **(G)**, and NAD(P)H measurements in Muc2-YFP-positive cells assigned as Paneth cells from their crypt location and morphology **(H)**, under the same treatment conditions as E,F. **I,J**: Corresponding images and phasor plots of villus regions **(I)** and NAD(P)H measurements in Muc2-YFP-positive cells assigned as goblet cells **(J)**. Measurements in H,J used Cellpose-derived reporter-positive cellular masks because reporter-associated crosstalk complicated TMRM-based mitochondrial selection. Masks were generated separately for each image and treatment condition. Quantitative summaries and existing comparisons are provided in Supplementary Table 14.

In organoids cultured for four days in mENR + CV, rotenone and oligomycin increased mitochondria-specific NAD(P)H intensity by 70.8% and 56.7%, respectively (Fig. 5A,B). Rotenone shortened τ(φ) from 1.733 to 1.381 ns and τ(m) from 3.249 to 2.907 ns. Oligomycin also shortened τ(φ), to 1.514 ns, whereas the decrease in τ(m) to 3.122 ns was not statistically significant. The IntestiCult-to-mENR protocol reproduced the intensity increases following rotenone and oligomycin treatments, by 50.4% and 45.5%, respectively (Fig. 5C,D). Rotenone shortened τ(φ) from 1.666 to 1.444 ns and τ(m) from 3.170 to 2.953 ns. Oligomycin produced a minor, non-significant reduction in τ(φ) to 1.598 ns. Its effect on τ(m) was also non-significant (3.207 ns). Collectively, rotenone produced more pronounced shortening of both τ(φ) and τ(m) of NAD(P)H than oligomycin. These differing lifetime responses suggested that the inhibitors alter the underlying fluorescence lifetime distribution in distinct ways, potentially reflecting how direct inhibition of NADH oxidation at complex I and inhibition of ATP synthesis affected this fluorescent cofactor pool (reduced form) and binding.

Combining NAD(P)H intensity with lifetime measurement can therefore help distinguishing effect of growth medium on the drug responses in future studies, capturing differences that intensity measurements alone cannot resolve. Notably, NAD(P)H-channel FLIM images of CV-supplemented organoids also contained bright, long-lifetime red–orange cytoplasmic puncta, which were most prominent under the IntestiCult + CV-to-mENR protocol and formed an additional long-lifetime phasor population consistent with oxidized-lipid-associated autofluorescence (Fig. 5A, C; 3× inset in C).

We next investigated the effects of drug treatment on the regions with active mitochondria in villus enterocytes (Fig. 5E,F): NAD(P)H intensity did not change from the control to rotenone, but increased non-significantly after antimycin A or oligomycin treatment. However, rotenone significantly shortened both τ(φ) from 1.679 to 1.451 ns and τ(m) from 2.970 to 2.737 ns, exactly as in stem cells (Fig. 5D). Antimycin A provided less drastic decrease of NAD(P)H τ(m) to 2.847 ns, non-significant with τ(φ) (1.548 ns). Strikingly, unlike in stem cells, oligomycin reduced enterocyte τ(φ) to a similar extent as rotenone (1.451 ns). However, oligomycin-induced τ(m) decreases followed similar patterns in enterocytes compared to stem cells.

We also took advantage of visualizing dynamics of NAD(P)H metabolism in Paneth and goblet cells, using the Muc2-YFP organoid line and assigning reporter-positive cells in crypt and villus regions as Paneth and goblet cells, respectively (Fig. 5G–J). Since the YFP spectral crosstalk complicated TMRM-based mitochondrial segmentation, we quantified NAD(P)H within reporter-derived whole cell masks. In crypt-residing Paneth cells, control NAD(P)H intensity averaged 14.99 photons/pixel, and none of the inhibitors significantly affected it (Fig. 5G,H). Rotenone treatment led to decrease of NAD(P)H τ(φ) from 1.563 to 1.329 ns and τ(m) from 2.884 to 2.686 ns, whereas Antimycin A did not show any effect. Interestingly, oligomycin showed either no effect or modest increase of NAD(P)H τ(m) This data indicates that in Paneth cells the ETC has lower activity than in stem cells and enterocytes, with NAD(P)H pool being mostly involved in non-OxPhos metabolic pathways such as glycolysis. In villus-confined goblet cells, the control NAD(P)H intensity averaged 19.20 photons/ pixel and remained statistically indistinguishable from control upon all treatments (Fig. 5I,J). However, the NAD(P)H lifetime was sensitive to rotenone (shortening of τ(φ) from 1.281 to 1.179 ns and τ(m) from 2.875 to 2.793 ns) and oligomycin (shortening of τ(φ), to 1.206 ns, but with non-significant τ(m) decrease). Antimycin A did not significantly reduce the NAD(P)H lifetime. Thus, both tested cell secretory populations showed reliable lifetime responses to complex I inhibition despite their weak NAD(P)H fluorescence.

We next looked if such a response pattern is maintained across multiple organoid lines. Thus, we looked at NAD(P)H-FLIM data for non-transfected organoid crypts (Fig. S13). Using TMRM to define the active mitochondria regions in crypts of non-labeled organoids, we found that both Antimycin A and Oligomycin significantly increased NAD(P)H intensity by ∼36%, contrasting with the minor (10%) increase induced by rotenone. Concomitantly, all the three inhibitors significantly shortened NAD(P)H lifetimes. Rotenone produced strongest decrease in τ(φ) and τ(m) (0.250 and 0.284 ns), followed by antimycin A (0.180 and 0.217 ns) and oligomycin (0.165 and 0.146 ns), reproducing the responses observed in Lgr5-GFP-labeled stem cells. For comparison, we measured TMRM-defined mitochondrial regions in YFP-negative cells in the crypts of Muc2-YFP organoids, observing similar trend in responses to Antimycin A, Oligomycin and rotenone, both in intensity and lifetimes (Fig. S13C). At the same time, villus regions without Muc2 expression in Muc2-YFP organoids showed results comparable to wild-type villus enterocytes (Fig. S13D and Fig. 5F).

Analysis of fluorescence lifetimes of NAD(P)H by decay fitting approach provided a complementary view of the responses presented in Figures 5B, D, F and H (Fig. S14; Supplementary Table 15): in the bi-exponential fitting data, rotenone decreased the mean short- and long-component lifetimes and increased the short-component amplitude fraction in all four datasets, consistent with the phasor lifetime shortening. Enterocytes showed a similar short-component lifetime upon rotenone and oligomycin treatment, but their long-component lifetime decreased after rotenone and remained close to control after oligomycin. The near-identical τ(φ) values after these treatments therefore arose from different component combinations. We also applied tri-exponential fitting since the bi-exponential high χ² and residuals values indicated that two components incompletely fitted the decays. The additional component resolved short, intermediate and long lifetime contributions whose amplitudes varied with treatment. Such heterogeneity is consistent with the multiple binding configurations described for NAD(P)H by Blacker and colleagues, ^16,17^ but assigning them to free, partially bound or fully bound cofactors would require additional molecular evidence.

Altogether, across the four major epithelial cell populations, rotenone consistently shortened NAD(P)H lifetimes, linking the observed fluorescence response to inhibition of complex I-dependent NADH oxidation. Responses to antimycin A and oligomycin varied with cell population and culture growth conditions. In stem-enriched regions, oligomycin produced smaller or statistically unresolved lifetime changes compared with rotenone; wild-type crypts showed the order of rotenone > antimycin A > oligomycin for both τ(φ) and τ(m) shortening. Enterocyte τ(m) followed such trend, whereas rotenone and oligomycin produced nearly identical τ(φ) decrease. The fitted component changes further highlighted such enterocyte responses. Together with the regional ATP measurements and transcriptomic signatures, observed patterns confirm that the relationship between ATP demand, electron transport and NADH oxidation varies across the crypt–villus axis.

### Modest NAD(P)H responses to metabolic pathways inhibition and regional differences in lipid-associated fluorescence

Having established cell type-dependent NAD(P)H responses to the drug treatment reflecting ETC and ATP synthase inhibition, we next looked how sensitive could the NAD(P)H responses be to the pathways supplying mitochondrial metabolism via pyruvate, glutamine and fatty acids. Thus, using wild type organoids, we measured regional NAD(P)H intensity, τ(φ) and τ(m), following treatment with UK-5099, targeting mitochondrial pyruvate uptake through the mitochondrial pyruvate carrier^66^, BPTES, targeting GLS1-mediated conversion of glutamine to glutamate^67^ and etomoxir targeting CPT1-dependent mitochondrial utilization of long-chain fatty acids^68^. Individual inhibitors produced modest, region-dependent reduction in NAD(P)H intensity, without significant changes in their phasor lifetimes (Fig. 6A,B). Combined application of the inhibitors produced specific but modest intensity reduction in crypt and villus regions (Fig. S15A,B). Overall, across all the treatment conditions, only the combined treatment showed significant lifetime shortening relative to control (Fig. S15). Crypt τ(φ) decreased from 1.647 to 1.552 ns, and τ(m) decreased from 3.094 to 2.992 ns. In villi, τ(m) decreased from 3.188 to 3.107 ns, whereas the τ(φ) decrease was nonsignificant (1.785 to 1.768 ns), which were smaller than those observed after rotenone in previous experiments. Collectively, the modest responses of NAD(P)H fluorescence could reflect the dynamic metabolic adaptation occurring in the multi-cell intestinal organoid model^58^, limiting the possibility to detect the effects of such inhibitors by only monitoring with NAD(P)H-FLIM method.

**Figure 6.**
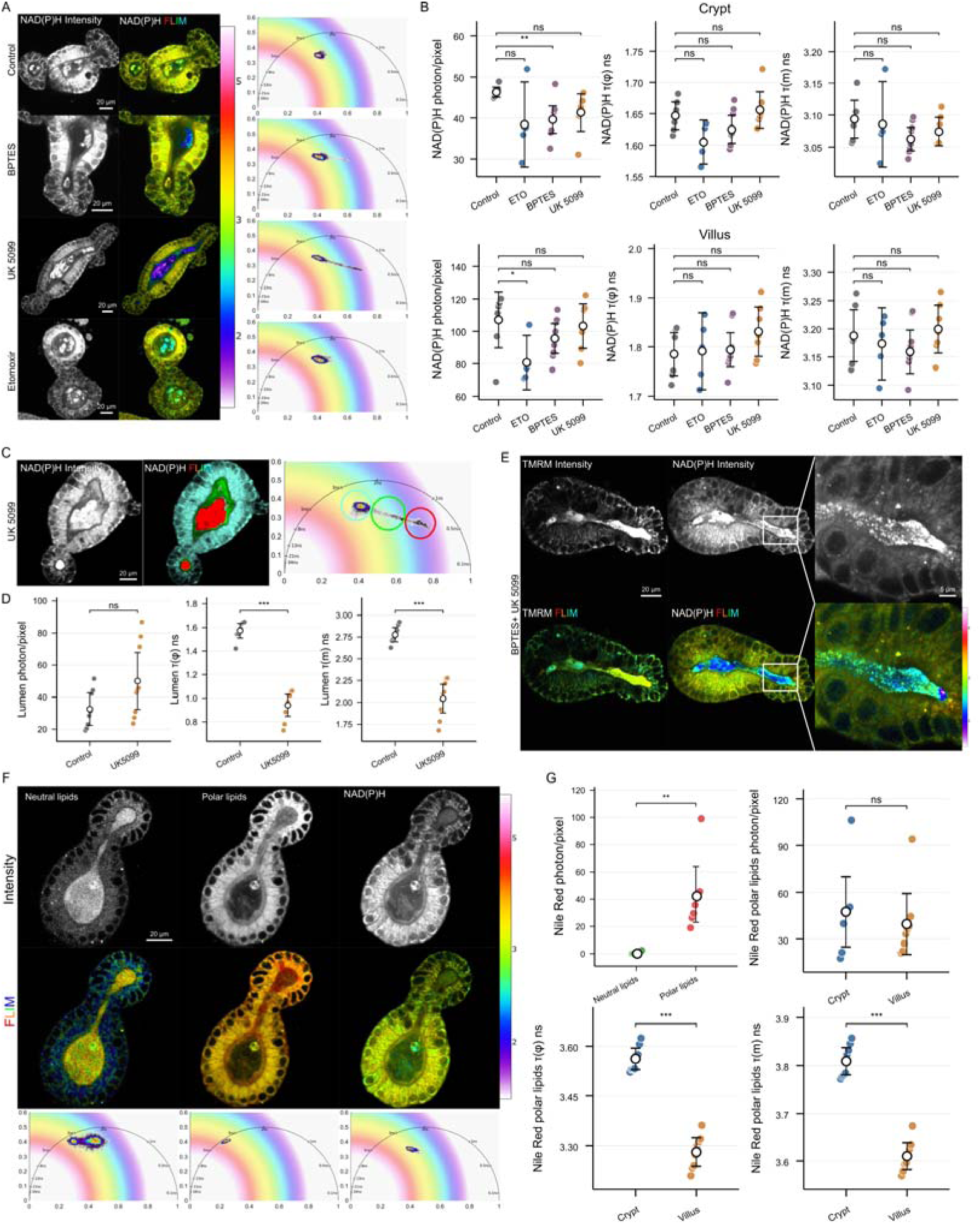
**Regional NAD(P)H responses to metabolic pathway inhibition and spatial variability in lipids. A,B**: Representative NAD(P)H intensity and FLIM images with corresponding phasor plots **(A)**, and quantification of NAD(P)H intensity, _τ_(_φ_), and _τ_(m), in crypt and villus regions **(B)**. Organoids were imaged under control conditions or following BPTES, UK-5099 or etomoxir treatment, n = 7 control, 5 etomoxir, 9 BPTES and 7 UK-5099 images per region. **C,D**: Representative NAD(P)H-channel intensity and phasor-selection-colored images following UK-5099 treatment **(C)**, and luminal fluorescence intensity, _τ_(_φ_) and _τ_(m) under control conditions and following UK-5099 treatment **(D)**. Red, green and cyan selections in the phasor plot map to debris-like luminal structures, other luminal contents and epithelial tissue, respectively; n = 8 control and 9 UK-5099 images. **E**: Representative TMRM and NAD(P)H intensity and FLIM images following combined BPTES and UK-5099 treatment. Enlarged regions show bright luminal granules near the crypt, with characteristically shorter lifetimes than in adjacent tissue (NAD(P)H channel). **F,G**: Representative Nile Red and NAD(P)H intensity and FLIM images with corresponding phasor plots **(F)**, and comparison of Nile Red emission-channel intensities and red-channel intensity, _τ_(_φ_) and _τ_(m) between crypt and villus regions **(G)**. Nile Red was excited at 488 nm and detected at 515–555 nm and 590–647 nm, preferentially sampling neutral-lipid-associated and more polar lipid environments, respectively. For the emission-channel comparison, crypt and villus mean intensities were averaged within each organoid; n = 8 paired organoid measurements. Quantitative summaries and existing comparisons are provided in Supplementary Table 16.

Interestingly, luminal organoid fluorescence showed a distinct response to UK-5099 in NAD(P)H spectral channel (Fig. 6C,D). Phasor FLIM mapped the shortest lifetime population to structures morphologically consistent with the luminal debris, with an intermediate population in the other luminal fluorescence species. Across the luminal measurements, τ(φ) decreased from 1.573 to 0.940 ns and τ(m) from 2.777 to 2.044 ns, while luminal intensity increased numerically from 32.55 to 50.05 units, without reaching significance. In addition, combined BPTES and UK-5099 treatment resulted in appearance of the bright luminal granules near the crypt regions. These granules exhibited shorter fluorescence lifetimes than the adjacent tissue in the NAD(P)H channel and were also strongly fluorescent on TMRM channel (Fig. 6E).

Both *de novo* fatty acid synthesis^69^ and fatty acid oxidation^70,71^ support Lgr5-positive intestinal stem-cell renewal and crypt maintenance in mice, motivating us to examine lipid-associated fluorescence along the crypt–villus axis. We therefore tested whether lipid imaging using solvatochromic fluorescent probes could complement our NAD(P)H-FLIM measurements. We performed staining with Nile Red, which shows both fluorescence intensity and lifetime dependence from the hydrophobicity of its environment^72–74^ (Fig. 6F,G).

Upon 488 nm excitation, this dye produces two spectral windows of fluorescence with 515–555 nm (neutral lipid-associated) and 590–647 nm (polar lipids)^72^. With live organoids, red fluorescence channel signals exceeded green intensity almost 20-fold also displaying drastically different lifetimes (Fig. 6F). Thus, we further focused on analysis of the polar lipids. We found that ‘red’ Nile Red emission was slightly brighter in crypts regions, compared to villi, while its lifetimes showed higher values in crypts: τ(φ), 3.562 versus 3.281 ns; τ(m), 3.809 versus 3.610 ns, indicating potentially different lipid composition across crypt-villus axis. To see if such heterogeneity was not an artifact, specific to Nile Red staining, we also performed staining with recently described solvatochromic dicarboxy-BODIPY BDP-DA1 dye^75^ (Fig. S15C,D): while we found difference intensity gradient across the crypt-villus, we also saw clear lifetime differences between these domains, again with longer lifetimes being visible in the crypt-based cells. Both tested dyes (Nile Red and BDP-DA1) have also detected basal membrane- and intracellular lipid droplets across the organoids, being more abundantly seen by BDP-DA1 staining. Collectively, our data indicates that the metabolic NAD(P)H gradient across the crypt-villus is also potentially connected with the lipid composition gradient.

## Discussion

### FLIM enables non-destructive multi-parameter imaging of cell metabolism in living organoids

We report successful application of the two-photon autofluorescence-based NAD(P)H-FLIM method combined with cell lineage-assisted labelling and complementary dye-based measurements of mitochondrial polarization, ATP, ROS and lipid-associated fluorescence, to examine cell-specific metabolism within living small intestinal organoids. Linking these readouts to epithelial cell position and applying metabolic perturbations extends the functional context of the optical metabolic imaging method.^8^ The intact organoid culture retains the spatial and temporal relationships between crypt-associated and differentiated cells that are normally lost in bulk or fixed cell measurements. For instance, time-lapse NAD(P)H-FLIM combined with TMRM-FLIM over 20 min upon oligomycin and tariquidar addition, helped to capture the dye redistribution alongside the distinct cellular NAD(P)H changes. Achieved combination of spatial and temporal information allows studying the acute responses within an organized epithelium and provides a foundation for extending metabolic imaging to dynamic three-dimensional measurements.

### NAD(P)H-FLIM enables constructing atlas of clearly distinct cell-specific metabolic phenotypes

Guided differentiation enriched epithelial populations with distinct NAD(P)H distributions while preserving variability among individual cells.^36^ Thus, membrane-based segmentation, live cell proliferation measurements, ACE2/MUC2 immunofluorescence and live reporters helped to attribute high NAD(P)H fluorescence to enterocytes and weak fluorescence to the goblet cells (Fig. 2). The stem–Paneth cells ‘metabolic contrast’ was less pronounced, although Muc2-YFP-positive crypt-residing cells displayed weaker NAD(P)H fluorescence than the neighboring stem cell-associated regions (Fig. 5). This pattern is consistent with the previously reported metabolic distinction between stem and Paneth cells.^22^ Comparing the four complete cell types describes the ‘metabolic’ phenotype produced by each culture condition. Future work can combine NAD(P)H intensity and lifetime with transmission light features, e.g. including Paneth- and goblet-cell granules, to build a live cell dynamic atlas of epithelial metabolic phenotypes.

However, our experiments with the metabolic pathway inhibitors, using e.g. UK-5099, BPTES and etomoxir (Fig. 6) have also revealed the ‘low sensitivity’ of NAD(P)H to the inhibition of the mitochondrial pyruvate transport, glutamine and fatty acids utilization. This data can be explained by the presence of compensatory mechanisms^58^ in living organoids, providing stable overall NAD(P)H levels and its involvement in cell bioenergetics. Yet they also indicate that the dynamics of NAD(P)H alone is not the only predictive indicator for the assessing complex effects on cell metabolism.

### Cell metabolic responses are highly dependent on the growth medium composition

Culture medium influenced both the epithelial lineage populations produced during differentiation and the resulting endogenous NAD(P)H fluorescence lifetimes at the time of imaging. Thus, ‘IntestiCult + CV’ growth conditions supported broader proliferation-associated Hoechst 33342 BrdU quenching and Lgr5-GFP labelling than the ‘mENR + CV’ growth medium. Organoids cultured in IntestiCult or mENR were imaged directly in their respective culture media. Only the imaging-medium group was switched shortly prior to the imaging, after culturing in mENR. The longer NAD(P)H lifetimes in IntestiCult growth medium therefore reflected influence of culture medium (Fig. S6C, D). Substrate supply could itself alter NAD(P)H binding and redox equilibria,^17^ providing a possible contribution to these medium-associated differences. The appearance of bright, long-lifetime puncta resembling lipid-droplet structures in the NAD(P)H channel of crypt zones under both mENR + CV and IntestiCult + CV-to-mENR conditions (Fig. 5A, C) may reflect medium-induced interference with mitochondrial oxidative and lipid metabolism, in part due to established effects of valproic acid on mitochondrial oxidative phosphorylation and fatty acid oxidation^76,77^. Moreover, the emergence of a characteristic long-lifetime shifted phasor signature consistent with oxidized lipids^78^ in stem-cell-enriched organoids following IntestiCult + CV induction (Fig. 5C) further suggests medium-dependent alterations in cellular redox state. However, since the full composition of IntestiCult is not publicly disclosed, the major components affecting intestinal cell metabolism in this culture remain unknown. Differences in the strength of inhibitor responses after mENR + CV culture and after the IntestiCult-to-mENR protocol further suggest that the ‘culture growth history’ can influence the subsequent metabolic response. Thus, future comparisons would benefit from the use of a common, well-defined imaging medium, matched exposure times and recorded growth histories. Testing defined medium components within the same organoid preparation would help separate immediate substrate responses from changes that develop during cell differentiation. Complementing NAD(P)H-FLIM of organoids with profiling of the NAD(P)H-binding enzymatic landscape (‘blueprint’) under matched growth conditions could further clarify metabolic trajectories during acute versus chronic pathway perturbations^79^.

### Metabolic crypt-villus gradient revealed in organoids

The crypt–villus axis showed strikingly opposing gradients in NAD(P)H fluorescence and mitochondrial polarization. Enterocyte-associated regions displayed higher NAD(P)H intensity and longer lifetimes, whereas TMRM labelling was stronger in crypts. Such relationship could reflect differences in fluorescent cofactor abundance,^10^ enzyme-binding configurations,^12,16,17^ mitochondrial organization,^23^ and the intracellular environment that influences fluorescence decay.^19^ The longer ATP-Red 1 lifetimes in villus regions were consistent with a larger ATP-related pool in enterocytes and aligned spatially with their higher NAD(P)H signal. Together, these readouts suggest that regional mitochondrial polarization and ATP utilization can vary in different directions across the crypt-villus axis.

Together, these spatial FLIM gradients reflect distinct, lineage-specific metabolic programs across the intestinal epithelium rather than a uniform cellular transition. Earlier work distinguished the metabolic identities of intestinal stem cells and neighboring Paneth cells and linked secretory differentiation to mitochondrial function and dynamics.^22,23^ Mitochondrial pyruvate transport also contributes to intestinal stem-cell maintenance and differentiation.^58^ Our spatial measurements extend this context to the enterocyte-associated high-NAD(P)H state and the weaker signals in goblet cells, which have different physiological demands. Linking these fluorescence phenotypes to cofactor abundance, redox balance and enzyme binding will help explain how they arise during differentiation.

To interpret the opposing TMRM and NAD(P)H gradients, we examined the contribution of apical dye transport. Efflux inhibition reduced luminal TMRM accumulation while preserving stronger crypt-associated labelling. We found that apical transport influenced dye distribution; nevertheless, a regional difference in TMRM retention remained after the efflux inhibition. CW-STED measurements complemented this result by revealing regional differences in apparent mitochondrial width and TMRM-positive skeleton length per tissue area. Together with the ATP measurements, these observations suggest that enterocytes can use the proton gradient more strongly for ATP synthesis, contributing to their lower steady-state polarization. The resulting polarization reflects the balance between proton pumping, ATP synthesis and proton leak.^53^

FLIM with ATP-Red 1 dye provided an ATP-sensitive readout alongside this polarization pattern. The dye lifetime increased with ATP concentration in solution, whereas organoid fluorescence intensity was also affected by mitochondrial polarization and treatment-associated redistribution. Lifetime measurements therefore helped to distinguish between ATP-related and probe accumulation changes. Longer baseline villus lifetimes and their shortening after oligomycin supported regional differences in ATP-related state and dependence on ATP synthase. Enterocyte-enriched TCA-cycle, respiratory-chain and ATP-synthase components expression in the public single-cell datasets provided further evidence for the observed regional transcriptional specialization in mitochondrial ATP production. Combining these expression profiles with the live imaging suggests a relationship between enterocyte metabolic organization and ATP demand that can be tested directly with paired functional measurements.

Respiratory inhibitors responses provided a complementary way to examine the link between ATP synthesis and NAD(P)H dynamics.^10,53^ The magnitude of inhibitor-induced τ(φ) shifts reflected how strongly mitochondrial NADH oxidation is coupled to ATP demand. In enterocytes, oligomycin produced a τ(φ) decrease comparable to rotenone, whereas Lgr5-GFP-enriched stem-cell regions and wild-type crypts showed higher sensitivity to rotenone than to the oligomycin. Rotenone directly restricts complex I-dependent NADH oxidation and oligomycin can suppress it indirectly by blocking ATP synthase and increasing the resistance to further proton pumping.^53^ Therefore, the enterocyte response was consistent with close coupling between ATP synthesis and NADH oxidation, as interrupting proton utilization perturbed the NAD(P)H lifetime almost as strongly as inhibiting complex I. The smaller oligomycin response in stem-enriched regions could reflect a lower ATP-linked contribution to NADH oxidation. This interpretation connects the observed response hierarchy to possible lineage-associated differences in oxidative phosphorylation.

The proposed difference in proton-gradient utilization also provided a framework for the regional ROS oxidation-sensitive signals. Higher crypt-associated ROS probe fluorescence coincided with stronger TMRM signals and a smaller observed ATP-Red 1 lifetime response to oligomycin. These observations suggest that the coupling between proton pumping and ATP-synthase activity differs between epithelial regions, allowing crypt-associated mitochondria to remain more polarized. Under appropriate respiratory and redox conditions, a high proton motive force can favor mitochondrial ROS formation.^80^ The crypt-associated highly polarized mitochondrial/ high ROS state could therefore contribute to redox signaling during epithelial renewal, e.g. for quick regulation of cell death during stem cell competition.^22^ Future more in-depth research is required for understanding the fine balance between mitochondrial NADH, mitochondrial polarization, ROS production and physiological O_2_ in the intestinal crypt-residing cells.

Regional lipid-associated fluorescence extended the crypt–villus differences beyond the NAD(P)H and mitochondrial readouts. Nile Red and BDP-DA1 showed longer lifetimes in crypt regions, consistent with differences in the local lipid environment sensed by these probes.^72,73,75^ Their intensity patterns differed, indicating that the two dyes sample lipid-associated structures and environments differently. Combining these measurements with targeted lipid analyses could connect the optical gradient to the membrane and lipid-droplet changes that accompany epithelial renewal.

### Apical transport and luminal fluorescence reveal compartment-specific responses

Here we also studied how the luminal fluorescence signals reflected the response to perturbations applied to the cells in organoids. Rhodamine 123, TMRM and ATP-Red 1 showed treatment-sensitive luminal accumulation, demonstrating how apical transport could influence the distribution of fluorescent probes in intact organoids. Oligomycin-associated depletion of luminal dye accompanied cellular mitochondrial responses. UK-5099 also shortened fluorescence lifetimes in the NAD(P)H detection channel within luminal material, despite the modest epithelial response. Combined UK-5099 and BPTES treatment produced bright luminal granules near crypt-associated luminal regions; these structures showed shorter NAD(P)H-channel lifetimes than the adjacent tissue and were also bright in the TMRM channel. The organoid lumen can therefore represent a distinct, treatment-sensitive fluorescent compartment. Measuring it separately preserves the epithelial metabolic readout while opening a route to studying how metabolism, secretion and cellular debris contribute to luminal contents and vice versa, even in the absence of functional microbiota.

## Conclusion

Combining NAD(P)H-FLIM with cell lineage assignment and complementary fluorescent readouts makes cell-associated metabolic differences accessible within the spatial organization of living intestinal organoids. The enterocyte-associated NAD(P)H, ATP-related signals, crypt-associated polarization, oxidation-sensitive signals and regional lipid fluorescence provide a framework for studying how epithelial functions are supported by different metabolic states. Three-dimensional time series, acquired under conditions evaluated for phototoxicity and stability, can further reveal metabolic changes preceding differentiation, regeneration or drug responses. Pairing these measurements with lineage validation, molecular profiling and targeted metabolic measurements can provide training data for fluorescence-based cell identification across organoid lines and culture batches. These applications will ultimately help connecting dynamic metabolic phenotypes to epithelial cell behavior during homeostasis and disease.

## Methods

### Organoid cultures

Mouse small intestinal organoids comprised the C57BL/6J and C57BL/6N lines and organoids carrying Lgr5-GFP or Muc2-YFP reporters. Wild-type organoids were established from *post mortem* mouse jejunal tissue kindly provided by Winter Vandenberghe and Noemi Castelluccio from the laboratory of Björn Heindryckx (Ghent University). Lgr5-GFP (Lgr5-EGFP-ires-CreERT2) mouse intestinal organoids^81^ were kindly provided by Prof. H. Clevers Lab (Hubrecht Institute, The Netherlands). Muc2-YFP intestinal organoids were obtained under material transfer agreement A21TT1005 (UGent–VIB, 2021). The reporter mice carry the Muc2-iCre driver and a Rosa26-YFP reporter^37^. The driver strain was C57BL/6J-Tg(Muc2-iCre)1Irc/Irc (MGI:8292155), carrying the Tg(Muc2-iCre)1Irc allele (MGI:8292154). Strain, sex, age, isolation date, special features and experiment assignments are provided in Supplementary Table 1. All experiments in this study were performed in vitro using previously established mouse intestinal organoid lines or organoids derived post-mortem from mouse jejunal tissue. In accordance with institutional guidelines at Ghent University and Belgian/European animal welfare regulations, post-mortem tissue collection solely for in vitro organoid culture is not classified as an animal experiment and therefore did not require an active Ethical Committee (EC) approval number. For newly established cultures, crypts were isolated from mouse jejunal tissue. Briefly, intestinal segments were flushed with ice-cold PBS, opened longitudinally, and cleared of mucus and villi. Tissue was cut into approximately 5-mm fragments and washed three times with cold PBS. Fragments were incubated in 2.5 mM EDTA for 10 min on ice, then transferred to fresh EDTA solution for 40 min at 4 °C. Crypts were released by vigorous shaking, passed through a 70-µm strainer, and collected at 300g for 5 min at 4 °C. After washing in Advanced DMEM/F12, approximately 200–400 crypts were embedded per 50-µl growth-factor-reduced Matrigel (356231, Corning) dome. Domes were polymerized for 5 min at 37 °C and overlaid with 500 µl mouse IntestiCult organoid growth medium (#06005, STEMCELL technologies) containing penicillin–streptomycin (15140122, Gibco).

Organoids were maintained in Matrigel with mouse IntestiCult medium or the indicated mENR formulation. Cultures were incubated at 37 °C, with 5% CO_2_. For passaging, Matrigel domes were disrupted mechanically, and organoids were fragmented by repeated pipetting. Fragments were collected at 300g for 5 min at 4 °C and re-embedded in fresh Matrigel. The basal medium and mENR formulation contained Advanced DMEM/F12, 10 mM HEPES, 2 mM GlutaMAX, 1× B27, 1× N2 and 1.25 mM N-acetyl-L-cysteine. Growth factors comprised EGF (50 ng / mL), Noggin (50-100 ng / mL) and R-spondin-1 (50-100 ng / mL). Reagent suppliers, catalogue numbers and final concentrations are listed in Supplementary Tables 2 and 3. For microscopy, organoids were seeded in 18-well glass-bottom chambers with # 1.5H coverslips (81817, Ibidi), using 2-5 µl Matrigel and 150 µl growth medium per well. Samples were equilibrated at 37 °C before imaging.

### Media, stem-cell enrichment and guided organoid differentiation

Stem-cell-enriched organoids were generated using IntestiCult supplemented with CHIR99021 (3 µM) and valproic acid (1.25 mM) mixture, designated as ‘CV’. For the guided organoid differentiation, we used method of Clevers et al^36^. Briefly, organoids were maintained in IntestiCult + CV for 2–3 days before transfer to mENR-based differentiation media. Paneth enrichment used mENR + CD, comprising CHIR99021 (5 µM) and DAPT (10 µM). Goblet enrichment used mENR + ID, comprising IWP-2 (5 µM) and DAPT (10 µM).

Enterocyte enrichment used mENR + I, comprising IWP-2 (5 µM). Differentiation was followed over 1–3 days as indicated.

For the Lgr5-GFP perturbation experiments in Fig. 5 the differentiation proceeded as follows: 1) 4 days in mENR + CV for Lgr5-GFP; 2) Two days in IntestiCult + CV followed by two days in mENR. For the medium comparison presented in Fig. S6C,D, organoids cultured in IntestiCult or mENR were imaged directly in their respective culture media. The imaging-medium (IM) experimental group was cultured in mENR and transferred shortly before acquisition to defined imaging medium consisting of phenol-red-free, bicarbonate-free DMEM (D5030, Sigma-Aldrich) supplemented with 10 mM D-glucose (G8270, Sigma-Aldrich), 2 mM GlutaMAX (35050-038, Gibco), 1 mM sodium pyruvate (11360-070, Gibco) and 10 mM HEPES-Na (pH 7.2; 15630-080, Gibco; Supplementary Table 2). Reagent suppliers, catalogue numbers and final concentrations are listed in Supplementary Table 4.

### Live organoid labeling with fluorescent dyes

Fluorescent dyes and tracers were diluted in basal medium for imaging. NR12S was used to label cell membranes^33^ and Zinpyr-1 was used to visualize Zn^2+^-rich granules in Paneth cells^38^. TMRM, ATP-Red 1, CellROX Deep Red and ROSstar 550 were used for mitochondrial membrane-potential-associated, ATP-associated and oxidation-sensitive fluorescence measurements. Rhodamine 123, Nile Red and BDP-DA1 were used in the corresponding transport and lipid-associated fluorescence experiments. Probe identities, stock solutions, staining concentrations, excitation/emission values, fluorescence lifetimes, suppliers, loading durations and washing conditions are summarized in Supplementary Table 5.

For labelling of cell proliferation with FLIM^29,35^, organoids were incubated with 100 µM BrdU for 18 h, followed by Hoechst 33342 staining for 30 min and 2-3 washes. Hoechst staining concentrations ranging from 50 nM to 2 µM were evaluated during optimization. Although the tracer table records a 200–400 nM working range, 400 nM was selected for subsequent combined Hoechst 33342/NAD(P)H imaging. BrdU-associated Hoechst quenching was assessed from nuclear fluorescence intensity and lifetime. Cellular NAD(P)H was quantified in separate NR12S-labelled samples, as described under cell and nuclear segmentation. Lgr5-GFP and Muc2-YFP reporter organoids provided complementary lineage information.

### Immunofluorescence

Whole-organoid immunofluorescence was adapted from Okkelman et al.^82^ Briefly, organoids were fixed in 4% paraformaldehyde for 10 min, permeabilized with 0.1% Triton X-100 for 10 min, and blocked in 5% FBS for 1 h at room temperature. Primary antibodies against ACE2 (PA5-47488, Invitrogen) and MUC2 (H-300, Santa Cruz) were diluted at 1:200 overnight at 4 °C. Samples were washed with TBST and incubated with Alexa Fluor 488 (A-11008, Invitrogen)/ Alexa Fluor 647 (A-21447, Invitrogen) secondary antibodies were diluted at 1:2000 for 1 h at room temperature, protected from light.

### Microscopy

Confocal and two-photon microscopy was performed on an inverted STELLARIS 8 Dive FALCON microscope (Leica Microsystems) equipped with a white-light laser (440−790 nm), a tunable infrared laser (IR Laser Mai Tai HP, 690-1040 nm), HC PL Apo 10×/0.4 air, HC Fluotar 25×/0.95 W, HC PL Apo 40×/1.25 GLYC corr, HC PL Apo 63×/1.4 oil objectives, 4Tune HyD NDD detectors, temperature / CO2-controlled incubator OKOLab (black, for light protection), and dedicated LAS X acquisition and analysis software (ver. 4.6.0 and ver. 4.8.1). Live organoids were imaged in glass-bottom chambers at 37 °C under the medium and environmental conditions specified for each experiment. Two-photon FLIM was used to measure endogenous NAD(P)H and combined Hoechst 33342/NAD(P)H fluorescence in BrdU-labelled organoids. Reporter fluorescence and fluorescent probes were imaged using the corresponding confocal FLIM acquisition settings in Supplementary Table 5. Recorded tracer-specific laser settings, scan speed, pixel dwell time, line accumulation and averaging, and frame accumulation and averaging are listed in Supplementary Table 5; two-photon acquisition combinations are listed in Supplementary Table 6.

**STED microscopy** to examine mitochondrial organization in TMRM-labelled live organoids was performed on STELLARIS 8 microscope equipped with an HC PL APO CS2 93×/1.30 NA glycerol-immersion objective with LAS X 4.9.0 (Leica Microsystems). TMRM was excited at 555 nm, and stimulated emission depletion was achieved using a continuous-wave 660-nm laser.

### Mitochondrial morphology analysis

The tissue-normalized morphology analysis used CW-STED images, comprising crypt samples and villus samples. We generated mitochondrial masks and skeletons with Nellie^54^, following its previous application to mitochondrial imaging in intestinal organoids^82^. We then quantified network morphology with custom Python scripts. Tissue boundaries were manually outlined on the TMRM images, excluding internal lumen and background spaces. Skeleton length was calculated by summing calibrated pixel links. Tissue-normalized skeleton density was the total skeleton length inside the tissue ROI divided by its physical area. Apparent local width was calculated at each skeleton pixel as twice the Euclidean distance to mask background. Each eligible non-edge component was summarized by its median local width. The field-level width was the median of component medians for components whose centroids lay inside the tissue ROI. Comparisons used tissue ROI -level, rather than individual components or skeleton pixels.

### Phasor FLIM data processing and data extraction

Intensity and phasor-coordinate images were exported from LAS X and imported into GSLab using the Leica/FALCON-compatible loading workflow (https://alexvallmitjana.github.io/GSLab). The recorded initial export used a threshold setting of 1 with filtering disabled and retained the G and S coordinates. GSLab was used for phasor visualization, masking and export of pixel- and object-level measurements. Background was removed by photon-intensity thresholding to remove pixels with photon counts below 5, and manually delineated lumen masks (Cellpose or LAS X) were excluded from epithelial measurements. Lumen masks were retained as separate measurement compartments for experiments targeting luminal fluorescence. Median filter and binning settings (downsampling in GSLab) were chosen based on specific processing workflow.

The phase lifetime, τ(φ), is derived from the phase delay of fluorescence emission relative to the excitation modulation. The modulation lifetime, τ(m), is derived from the attenuation of the emission modulation. The two agree for a single exponential decay and can differ when several decay components contribute to the signal. For phasor coordinates G and S at angular frequency ω, the phase and modulation lifetimes are defined as 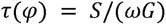 and 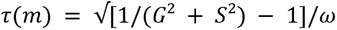. Here, ω = 2πnf, with f defined by acquisition frequency and n defined by harmonic. Pixel exports contained coordinates, intensity, G, S, τ(φ), and τ(m) available in selected datasets. Image-region (crypt region or villus region) measurements were arithmetic means of exported pixel values within corresponding masks that manually delineated in Cellpose. GSLab-exported CSV measurements were analyzed using custom Python scripts, with final masks applied where required.

### Fluorescence lifetime decay fitting

Native NAD(P)H photon histograms were extracted from matched Leica acquisitions and summed within registered cellular masks for Figures 1 and 2 or within the image-specific measurement ROIs for Figure 5. Each cellular or image ROI was fitted independently. The model was a sum of two or three exponential decays, re-convoluted with the acquisition-matched calculated instrument response function (IRF). The sampled IRF shape was retained, with a fitted temporal shift and background. The model integrated exponential emission across native photon-bin widths and accounted for preceding excitation pulses. Fits minimized Poisson deviance using bounded nonlinear least squares and multiple starting values; the finite solution with the lowest deviance was retained. The terminal clamped/overflow bin was excluded from the fitting objective. Lifetimes were bounded between 0.005 and 20 ns and ordered as τ < τ < τ where applicable. Figures 1 and 2 use bi-exponential fits; Figure 5 includes bi- and tri-exponential fits. Relative pre-exponential amplitudes were α = 100A /ΣA . Amplitude-weighted and intensity-weighted lifetimes were calculated as τA = ΣA τ /ΣA and τI = ΣA τ ²/ΣA τ, respectively. Fitted component and weighted lifetimes describe the reconvolved decay, whereas τ(φ) and τ(m) are calculated from phasor coordinates.

### Cell and nuclear segmentation

NR12S membrane images were segmented in Cellpose. Masks were manually corrected to remove lumen debris and detached objects outside the organoid epithelium. Corrected masks were applied to the corresponding FLIM images in GSLab, and object-level intensity, lifetimes and centroid coordinates were exported.

For nuclear analysis, Hoechst FLIM images were rendered using τ(m) magnitude combined with intensity to improve visualization of quenched nuclei in Cellpose. Nuclear masks were manually corrected to exclude luminal and detached nuclei before object-level measurements were exported. The Hoechst-associated phasor population persisted after nuclear exclusion in the combined-staining dataset. Thus, principal cellular NAD(P)H measurements therefore used the separate NR12S/NAD(P)H acquisitions. For the displayed differentiation summaries, values outside Q1 − 3×IQR to Q3 + 3×IQR were excluded separately within each condition and metric.

### NAD(P)H photon-count optimization

Eight acquisition settings were screened and grouped into low-, intermediate- and high-photon regimes (Supplementary Table 6). Organoid-level intensity and lifetime measurements were compared across settings, and zoom was evaluated separately. NR12S-derived cell masks were used to measure the availability of cellular phasor lifetimes at different photon counts. Background thresholding retained pixels above the recorded photon-count cutoff (5 photons/pixel in the general workflow; 5–10 in the Figure S1 examples), and the phasor selection was restricted to the recorded G and S ranges, 0–1 and 0–0.5. For each metric, a cell was counted as valid when its exported ROI value was finite and calculable, and invalid when that value was absent or non-finite. Thus, ‘valid ROI’ describes lifetime availability after processing. No separate numerical minimum-pixel threshold was documented for this classification; ‘insufficient’ refers to a missing calculable ROI value. All photon-count groups were analysed with median-filter setting 5 and no downsampling. Low-photon images were additionally processed with median-filter settings 5, 10, 20 and 40 and image dimensions of 512 × 512, 256 × 256 and 128 × 128 pixels. Intensity, τ(φ), τ(m) and valid-ROI fractions were compared across these processing conditions.

### Spatial analysis along the crypt–villus axis

For cell-based spatial analysis, exported cellular centroids were analyzed using a custom MATLAB script. Crypt-base and villus-side anchor cells were selected interactively for each sample. Centroids were projected onto the axis connecting these anchors, and positions were normalized from 0 at the crypt side to 1 at the villus side. Measurements were grouped into 50 equal-width bins for spatial summaries.

The TMRM/NAD(P)H profiles in Figure 3 used the separate grouped pixel-export workflow. These profiles pooled observations into 50 bins, centered from 0.01 to 0.99, without additional smoothing. Where displayed, intervals were calculated as the pooled mean ± 1.96 SEM.

### Rhodamine 123 and TMRM efflux inhibition assays

Reagents used for metabolic and transport perturbations are listed in Supplementary Table 7, with their reported functions, stock and working concentrations, solvents, suppliers and catalogue numbers. Experiment-specific concentrations and treatment combinations are described in the corresponding assay sections below.

Rhodamine 123 was first used in preliminary luminal-transport assay. Treatments included tariquidar (5 µM) or oligomycin (10 µM). Luminal intensity was summarized per image, and crypt-adjacent and villus-adjacent lumen regions were paired within acquisitions where available. TMRM was maintained at 10 nM in the imaging medium during the efflux experiments. The initial mitochondrial-perturbation screen used 2 µM FCCP, 10 µM rotenone or 10 µM oligomycin. Oligomycin concentration-dependent response experiments employed 100 nM, 500 nM, 1 µM and 10 µM. Tariquidar and elacridar series used 100 nM, 500 nM, 1 µM and 5 µM. For each condition, regional TMRM and NAD(P)H intensity and lifetimes were extracted from the corresponding masks. Paired crypt-minus-villus TMRM differences were calculated for acquisitions containing both regions. Luminal fluorescence was measured separately from epithelial fluorescence. Time-lapse TMRM/NAD(P)H acquisitions included a baseline frame followed by tariquidar at 500 nM or oligomycin at 10 µM. Ten frames, t0–t9, were analyzed per series. For each trajectory and channel, intensity was normalized to its own baseline as F(t)/F(0). Individual trajectories and timepoint means were displayed, with t-based 95% confidence intervals across trajectories.

### ATP-Red 1 dye calibration and regional analysis

We calibrated 5 µM ATP-Red 1 at ATP concentrations of 0, 1, 5, 10, 20, 40, 60 and 80 mM in 25 mM HEPES-Na, pH 7.8. For each calibration image, pixels were retained when all required intensity, phasor and lifetime values were finite. Per-image intensity and lifetime values were calculated as arithmetic pixel means and summarized across the five images per concentration, whereas phasor trajectories were calculated across the five image centroids with SEM crosshairs. Acquisition settings and inferred spatial binning were checked for consistency across the calibration series. Calibration trajectories were descriptive and were not used to assign absolute intracellular ATP concentrations.

Organoids were labelled with ATP-Red 1 (5 µM), as summarized in Supplementary Table 5, and imaged together with endogenous NAD(P)H. Crypt and villus regions were paired by acquisition for the control comparison. Treatments comprised tariquidar at 500 nM, oligomycin at 10 µM and rotenone at 10 µM.

### ROS probe assays and regional analysis

CellROX Deep Red and ROSstar 550 dyes were used to measure oxidation-sensitive fluorescence in crypt and villus-like regions. CellROX was imaged alone in the control dataset and with TMRM and NAD(P)H in the inhibitor-associated multichannel dataset. Tariquidar or elacridar was used at 500 nM in the indicated experiments. ROSstar 550 was examined under control conditions and during 500 nM tariquidar treatment. Regional comparisons were paired within each image. Analyses compared crypt and villus regions within each condition.

### Metabolic pathway perturbations

Organoids were treated with UK-5099, BPTES or etomoxir at 10 μM individually, with all three pairwise combinations, or with the three inhibitors together. Epithelial NAD(P)H intensity, τ(φ) and τ(m) were summarized separately for crypt and villus image regions. For UK-5099 luminal experiment, lumen-associated fluorescence was measured using dedicated regional exports. Phasor selections were mapped back to the images to visualize spatially distinct luminal populations. Treatment-associated puncta were described from their fluorescence distribution and morphology.

### Lipid imaging with Nile Red and BDP-DA1 dyes

Nile Red fluorescence was analysed in the two emission channels, representing neutral lipids (515–555 nm) and polar lipids (590–647 nm). For neutral lipids-versus-polar lipids comparison, crypt and villus regional means were weighted equally within each organoid. Crypt and villus intensity and lifetimes analyses used common valid exported coordinates in polar lipids channel. BDP-DA1 regional intensity and both lifetime metrics were compared within matched image pairs.

### Lgr5-GFP-defined compartments and mitochondrial NAD(P)H

For Lgr5-GFP datasets, GFP, TMRM and NAD(P)H measurements were aligned using common exported pixel coordinates. Image-specific GFP intensity percentiles defined low, intermediate and high compartments; intermediate and high compartments were combined into a GFP-enriched selection representing approximately 75% of valid pixels. TMRM-derived mitochondrial masks were generated with an infer-subc-style workflow.^83^ Within the common valid domain, TMRM intensity was normalized between its 1st and

99.8th percentiles and clipped to 0–1. A 3-pixel median filter and Gaussian smoothing (σ = 1.34 pixels) preceded detection of punctate structures with a negative Laplacian-of-Gaussian response (σ = 1.5 pixels) and filamentous structures with a Sato filter (σ = 1 pixel). Each response was normalized between its 1st and 99.5th percentiles. Pixels passing either the dot-response threshold of 0.05 or filament-response threshold of 0.11 formed the initial mask. Connected objects smaller than 12 pixels were removed, holes smaller than 12 pixels were filled, and binary closing used a disk of radius 1 pixel. The final mask was restricted to valid coordinates and intersected with the GFP-enriched selection for NAD(P)H quantification. The same TMRM segmentation procedure was used for wild-type crypt and villus measurements.

Masks were generated independently for each control and treated image, so they identify TMRM-retaining structures under the corresponding condition. Parameters and implementations are available in the analysis repository.

### Muc2 reporter segmentation

Muc2-YFP cellular masks were generated using Cellpose-SAM^84^. Objects with reporter intensity at or above the image-specific 75th percentile formed the reporter-positive mask; remaining valid pixels defined the reporter-negative compartment. Reporter-positive crypt and villus regions were assigned as Paneth- and goblet-cell regions, respectively. NAD(P)H was quantified across reporter-positive cellular masks because possible YFP contribution to TMRM detection complicated mitochondrial selection. Reporter-negative mitochondrial masks were generated separately from TMRM images.

Detailed segmentation and filtering parameters are specified in GitHub.

### Public database transcriptome analysis for single-cell RNA-sequencing data

Public mouse intestinal organoid datasets GSE164638^42^ and GSE165092^43^ were reanalyzed using Scanpy. The untreated ENR subset comprised 18,806 cells from five source-sample groups; pooled dome cultures were treated as one group. Processing included sample-aware quality control, scDblFinder doublet removal, normalization, log transformation and Harmony integration. TCA-cycle, respiratory-chain and nuclear ATP-synthase signatures were assessed in enterocytes, stem cells, goblet cells and Paneth cells. Scores were averaged within each source group and cell type, then combined with equal weighting across source groups. Expression summaries were descriptive. Detailed preprocessing and annotation parameters, gene sets and software versions are specified in GitHub.

### Statistical analysis

Regional FLIM analyses used image-level compartment means as the analysis unit, with matched crypt–villus measurements paired by acquisition. Unless otherwise specified, endpoint plots show individual values and means with t-based 95% confidence intervals. Multigroup endpoint comparisons followed Shapiro–Wilk normality testing within each condition and metric. When all groups passed, analyses used Welch ANOVA and Games–Howell-type comparisons against control. Otherwise, Kruskal–Wallis tests and Dunn comparisons with Holm correction were used. Paired contrasts used one-sample t-tests on within-image differences or Wilcoxon signed-rank tests, according to normality. Independent two-group comparisons used Welch t-tests or Mann–Whitney U tests.

Figure-specific exceptions, correction families, sample counts, effect estimates and exact P values are documented in the statistical outputs and analysis code. Significance labels refer to adjusted P values where multiple-comparison correction was applied and unadjusted P values otherwise: *, P < 0.05; **, P < 0.01; ***, P < 0.001, ****, P < 0.0001; ns, not significant.

## Software and data availability

Image acquisition and processing used LAS X (4.6.0 and 4.8.1), GSLab (1.289), Cellpose (4.0.1), Nellie (1.0.4), MATLAB (2024a) and custom Python scripts. Python analyses used NumPy (2.2.6), pandas (2.3.3), SciPy (1.15.3), scikit-image (0.25.2) and Matplotlib (3.10.9); the reproducible analysis environment used Python 3.10.9. The Zenodo deposit (DOI 10.5281/zenodo.23103396) contains the microscopy data package, including microscope-exported TIFFs and XML metadata, GSLab-generated images and CSV measurements, masks, source-data tables and statistical outputs. Public transcriptomic datasets are identified by GSE164638 and GSE165092. Two accompanying Excel workbooks contain the numbered supplementary tables: “Supplementary Tables 1–7 — Materials and experimental settings.xlsx” and “Supplementary Tables 8–16 — Quantitative results and fitting data.xlsx”. Each numbered worksheet contains one supplementary table.

Analysis parameters, statistical procedures and figure-specific workflows are documented in the public GitHub repository (https://github.com/HangZhouFLIM/organoid-flim-analysis). The repository includes software dependencies, input-data mappings and instructions defining the scope of each reproducible workflow.

## Use of Artificial Intelligence

OpenAI Codex was used to assist with drafting initial manuscript as well as development, debugging and documentation of the analysis code. The authors reviewed the AI-assisted manuscript text and analysis code, together with statistical outputs and plots. The authors take full responsibility for the accuracy and integrity of the analyses, interpretations and final manuscript.

## Supporting information

Experimental Settings

Experimental Supplementary Tables

## Acknowledgements

We are grateful to A. Vallmitjana for support with GSLab software, G. Zanetti for providing CellROX Deep Red, M. Filatov for sharing BDP-DA1, N. Castelluccio and W. Vandenberghe for providing mouse jejunal tissue for wild-type organoid cultures.

This work was supported by the Special Research Fund (BOF) grants (BOF/STA/202009/003, BOF/BAF/1 y/25/1/004, BOF/BAF/4y/26/1/004 and BOF.ITN.2026.0010.01), Research Foundation Flanders (FWO, I001922N, I004124N), and the European Union, fliMAGIN3D-DN Horizon Europe-MSCA-DN No. 101073507 grants.

## Author Contributions

H.Z. designed and performed the experiments, established and optimized the two-photon NAD(P)H-FLIM acquisition and multiplexed live-organoid imaging workflows, carried out organoid culture, pharmacological perturbations, immunofluorescence validation, and ATP-Red 1 calibration, developed and executed the image-segmentation, phasor-analysis, decay-fitting, and statistical pipelines, analyzed and visualized the data, and wrote the original draft of the manuscript. I.A.O. contributed to experimental design and planning, established and maintained organoid cultures, assisted with data interpretation, and revised the manuscript.

A.D.K. performed organoid experiments and assisted with data acquisition. L.L. performed reanalysis of the publicly available single-cell RNA-sequencing datasets. L.V. and B.N.L. provided the transgenic Muc2-YFP intestinal organoid line and contributed to the manuscript.

K.J. provided access to the FLIM–STED microscopy, contributed to the experimental design of the ATP-Red 1 calibration, and critically reviewed the manuscript. R.I.D. conceived and supervised the project, secured funding, contributed to experimental design and data interpretation, reviewed and edited the manuscript. All authors read and approved the final manuscript.

## Competing Interests

The authors declare no competing interests.

## Notes

https://github.com/HangZhouFLIM/organoid-flim-analysis

### Competing Interest Statement

The authors have declared no competing interest.

