## Supplementary Figures S1-S15 for "NAD(P)H-FLIM reveals spatial and temporal epithelial metabolic heterogeneity in live intestinal organoids"

Supplementary tables are supplied in separate Excel workbooks.
Supplementary Tables 1–7 — Materials and experimental settings.xlsx
Supplementary Tables 8–16 — Quantitative results and fitting data.xlsx

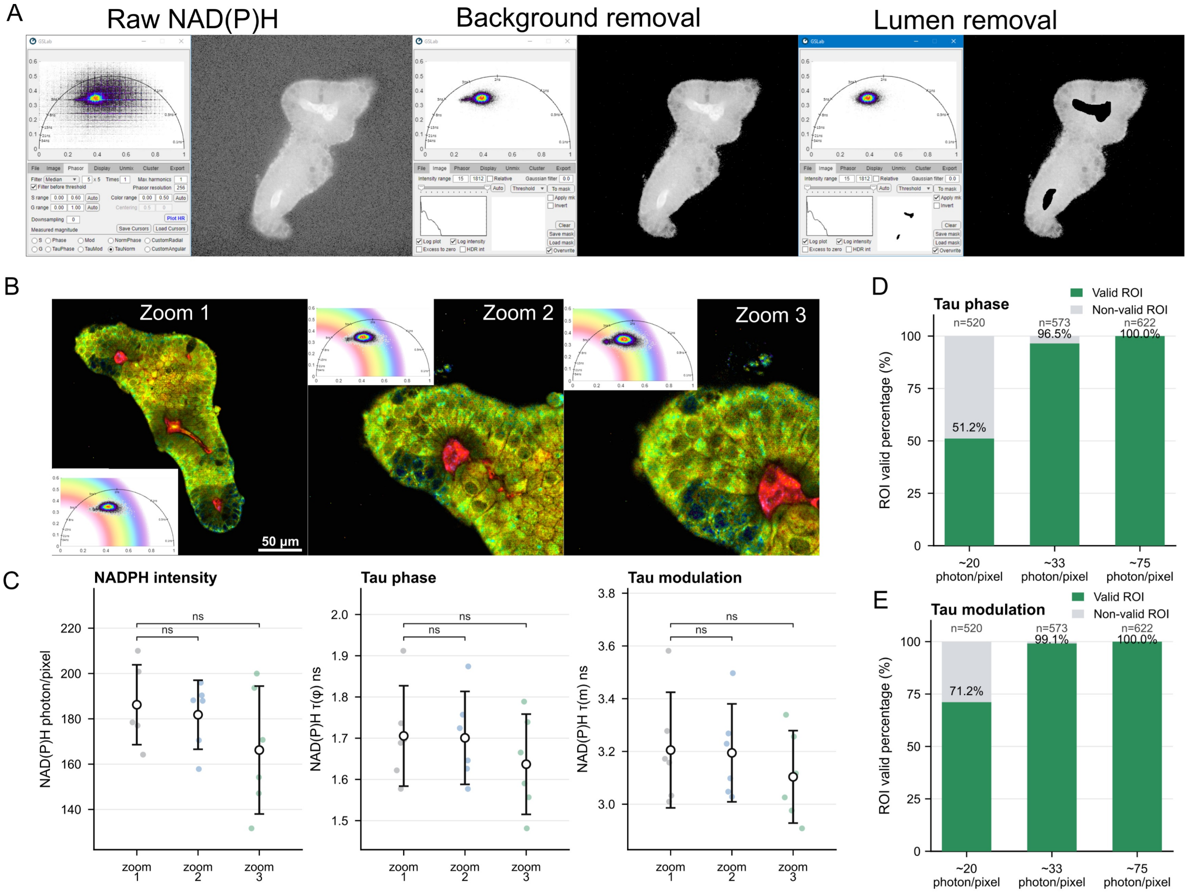


**Supplementary Figure S1. NAD(P)H phasor-FLIM processing and lifetime analysis pipeline with intestinal organoids. A**: Organoid-level NAD(P)H signal processing workflow. Background pixels were removed using a photon-count threshold, photon counts <5–10. Lumen-associated signal was excluded by applying manually drawn lumen masks in GSLab followed by mask inversion. For single-cell analysis, cell masks generated from Cellpose. Phasor plots show pixel distributions from raw NAD(P)H images, background-removed images, and lumen-removed images. Details in Methods, 'Phasor analysis' and 'NAD(P)H photon-count optimization'. **B**: Representative organoid NAD(P)H-FLIM images acquired from zoom 1 to zoom 3 and visualized by phasor-based pseudocoloring. Scale bar: 50 µm. Phasor plots: median filter 5. **C**: Average NAD(P)H photons/pixel, τ(φ), and τ(m) from zoom 1 to zoom 3. Six samples were analyzed per group. **D**: Percentage of cells with valid and invalid τ ROIs under ~ 20 photons/pixel, 33 photons/pixel, and 75 photons/pixel in intestinal organoids. A total of 520, 572, and 622 cells were analyzed, respectively. Both τ(φ) and τ(m) were evaluated. Valid ROIs have a finite exported lifetime value; invalid ROIs lack a calculable value after pixel selection. The operational criteria are described in Methods, 'NAD(P)H photon-count optimization'.

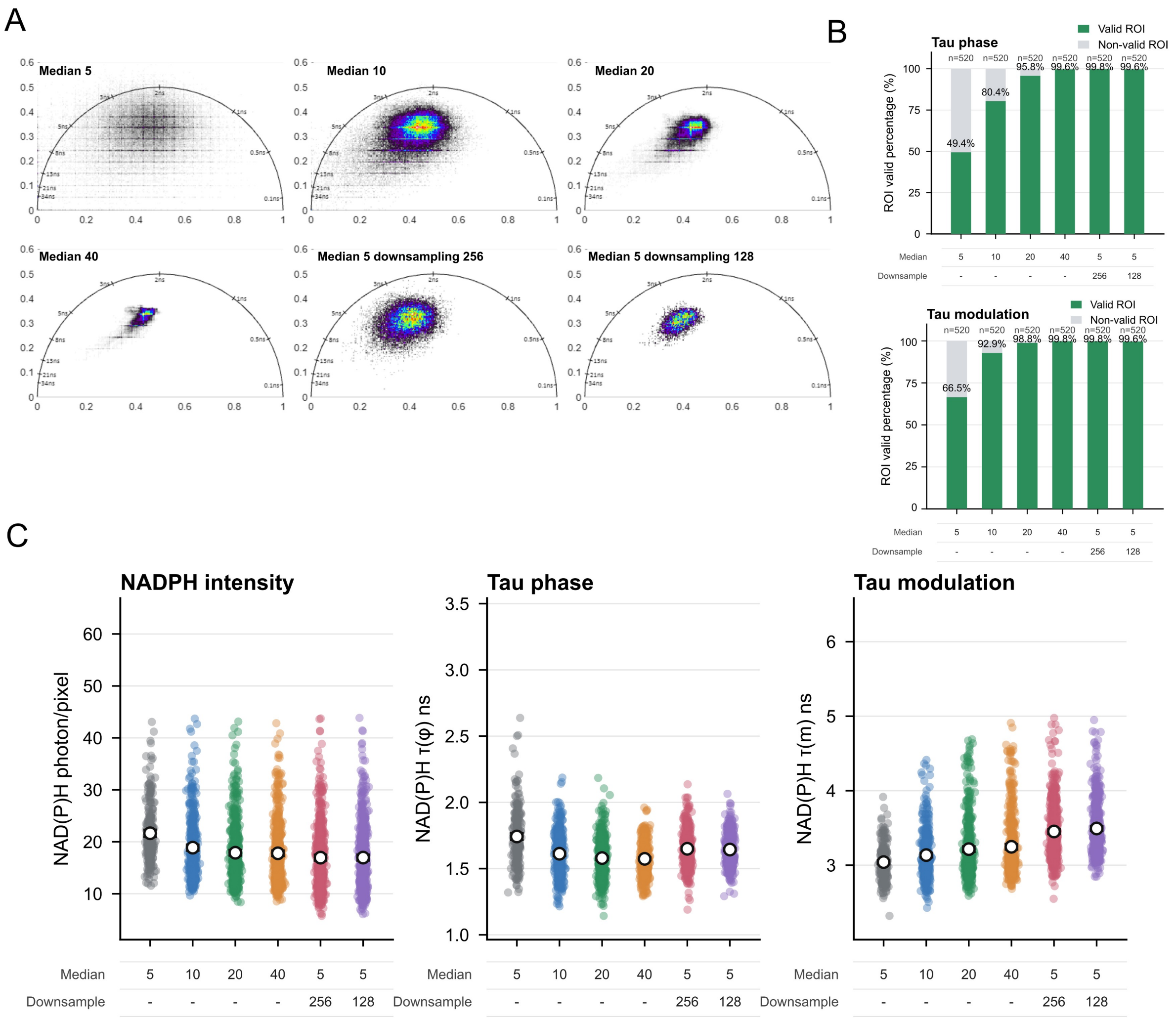


**Supplementary Figure S2. Effect of median filtering and downsampling on phasor-FLIM analysis under low-photon conditions.**

**A**: Phasor plots obtained under ~ 20 photons/pixel condition with different median filter and downsampling settings. Median filter sizes of 5, 10, 20, and 40 were applied, and images were downsampled from the original 512 × 512 pixels to 256 × 256 or 128 × 128 pixels, similar to the binning function in LAS X FLIM. **B**: Percentage of cells with valid or invalid ROIs under after adjustment of median filtering and downsampling parameters. In total, 520 cells were analyzed. **C**: Single-cell mean intensity and the corresponding average NAD(P)H τ(φ) and τ(m) values after applying different median filter and downsampling settings under approximately 20 photons/pixel conditions.

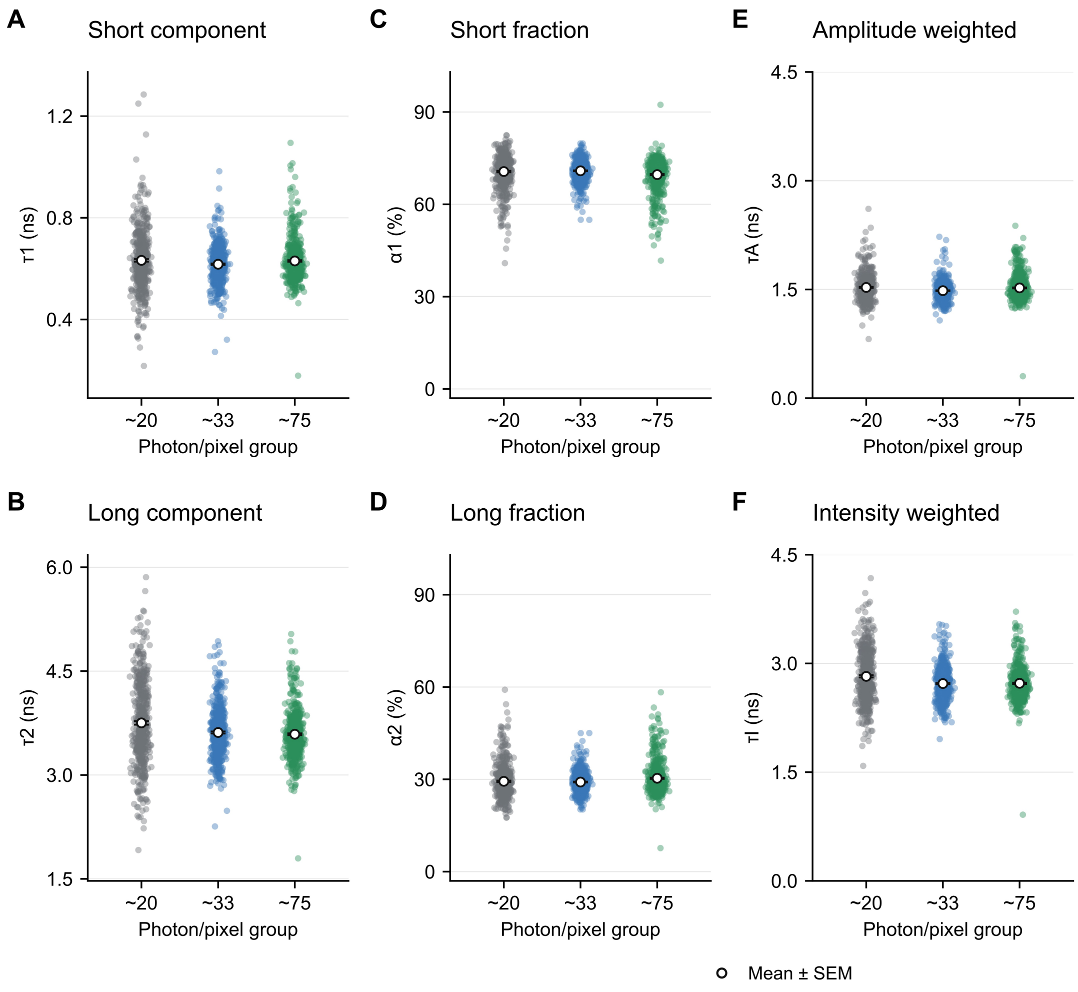


**Supplementary Figure S3. Fitting decay (double-exponential fit) analysis of cellular NAD(P)H fluorescence lifetimes for experiments presented in Fig. 1. A,B**: Short- and long-component lifetimes ($\tau_1$ and $\tau_2$). **C,D**: Corresponding amplitude fractions ($\alpha_1$ and $\alpha_2$). **E,F**: Amplitude-weighted (τA) and intensity-weighted (τI) lifetimes. Each point represents one independently fitted cell; white markers and error bars show the cell mean ± SEM. The approximately 20, 33 and 75 photons/pixel groups contain 520, 573 and 622 cells, respectively, from four acquisitions per group. All saved fitted values are included. Component definitions and reconvolution procedures are described in Methods, 'Native fluorescence-decay fitting'; summaries are in Supplementary Table 9.

A

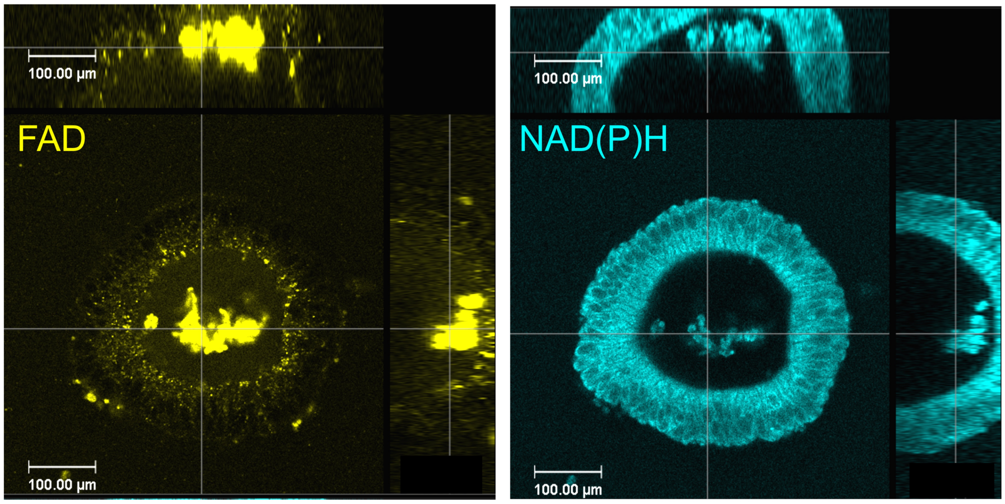


B

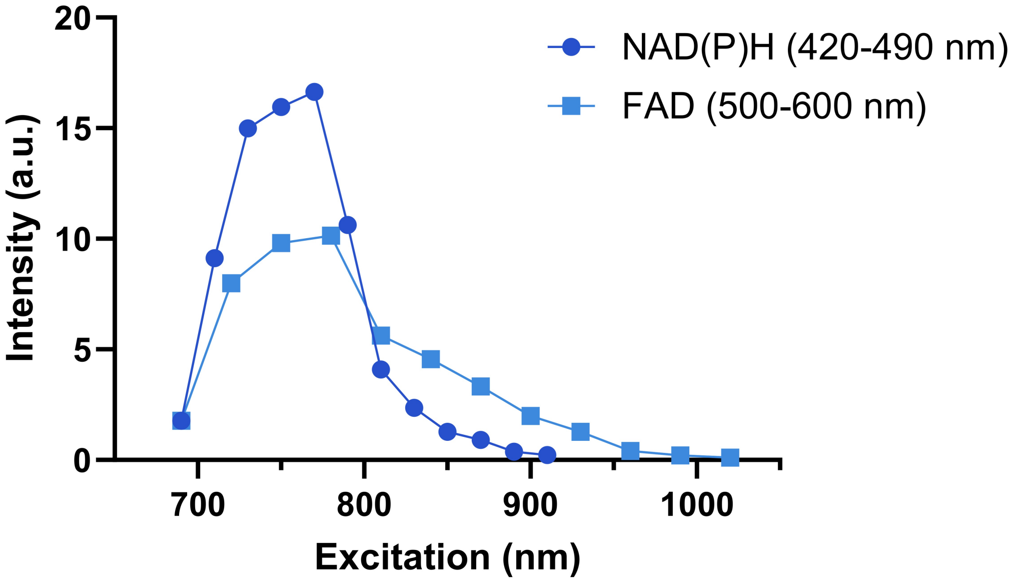


**Supplementary Figure S4. Autofluorescence imaging and excitation spectra of NAD(P)H and FAD in a porcine intestinal organoid.**

**A**: Orthogonal 3D views of endogenous FAD and NAD(P)H fluorescence in a live porcine small intestinal organoid. FAD was imaged using 890 nm excitation, 12% laser power, 400 Hz scan speed, 2.8 µs pixel dwell time, and 3-line accumulation. NAD(P)H was imaged using 740 nm excitation, 8% laser power, 400 Hz scan speed, 2.8 µs pixel dwell time, and 2-line accumulation. **B**: Excitation spectra in a porcine intestinal organoid. Mean fluorescence intensity (a.u.) was recorded in the 420–490 nm detection window as excitation was varied from 690 to 910 nm, and in the 500–600 nm window as excitation was varied from 690 to 1020 nm.

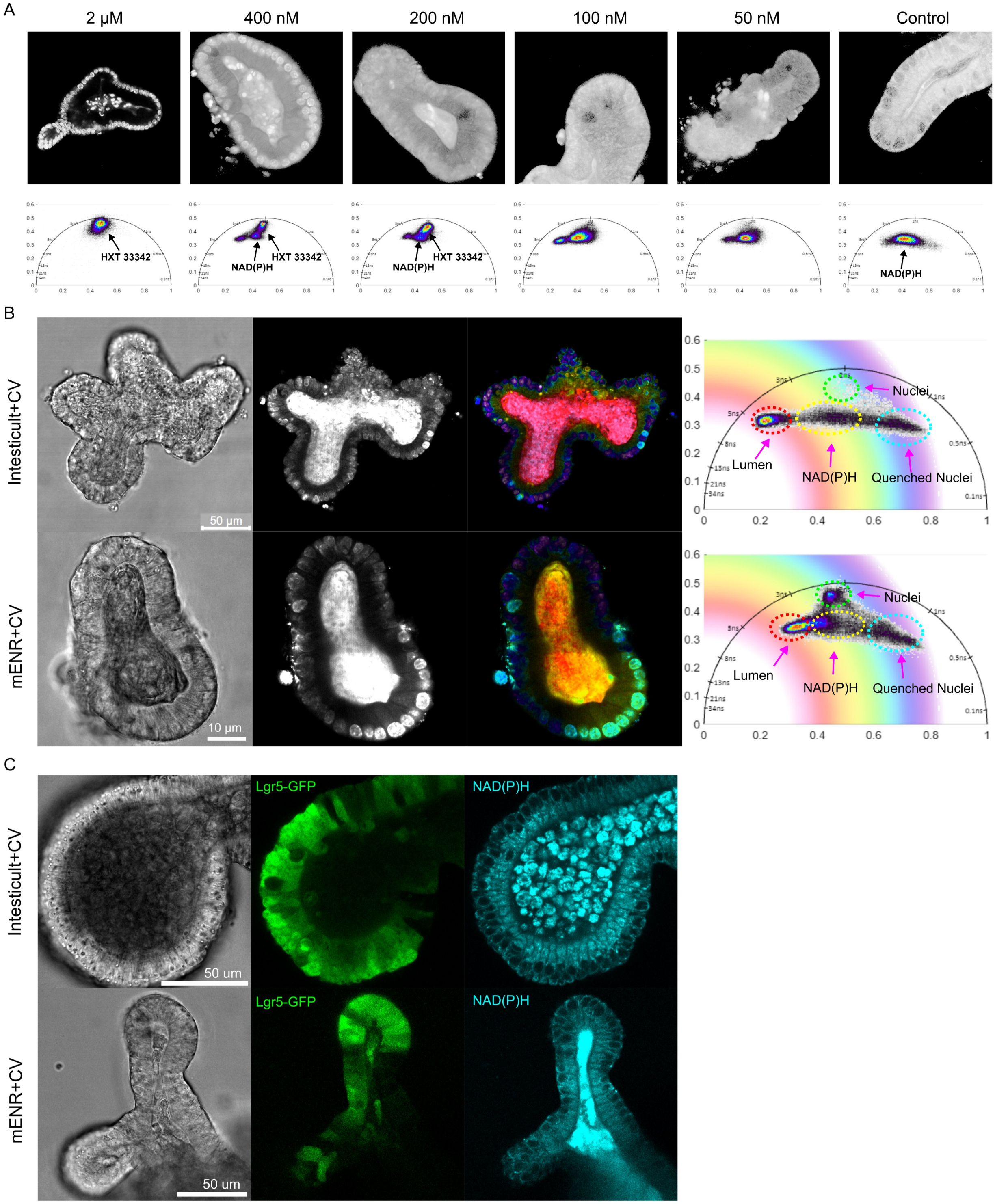


**Supplementary Figure S5. Optimization of Hoechst 33342 concentration and validation of stem cell-enriched culture conditions for BrdU–Hoechst 33342 FLIM/ NAD(P)H FLIM imaging.**

**A**: Optimization of Hoechst 33342 concentration for simultaneous visualization of nuclear Hoechst 33342 and cellular NAD(P)H signals in the NAD(P)H emission channel, 410–490 nm. Hoechst 33342 was tested from 50 nM to 2 μM, and the corresponding phasor plots show the lifetime separation of Hoechst 33342 and NAD(P)H signals. nuclear Hoechst 33342 signal was bright and obscured the cellular NAD(P)H signal reading at 2 μM. nuclear signal was negligible at 50–100 nM. the. Both 200 and 400 nM enabled detection of nuclear Hoechst 33342 and cellular

NAD(P)H signals. **B**: Comparison of stem cell-enriched organoids cultured in commercial IntestiCult + CV medium or published mENR + CV medium using BrdU–Hoechst 33342/ NAD(P)H FLIM imaging. Nearly all nuclei (dark purple) were Hoechst-quenched in IntestiCult + CV, a subset of nuclei (bright cyan) remained unquenched in mENR + CV. Phasor plots distinguish lifetime signals from the lumen, cellular NAD(P)H, unquenched nuclei, and quenched nuclei. **C**: Validation of stem cell enrichment in IntestiCult + CV and mENR + CV conditions using Lgr5-GFP organoids. Both Lgr5-GFP and NAD(P)H intensity signals were detected.

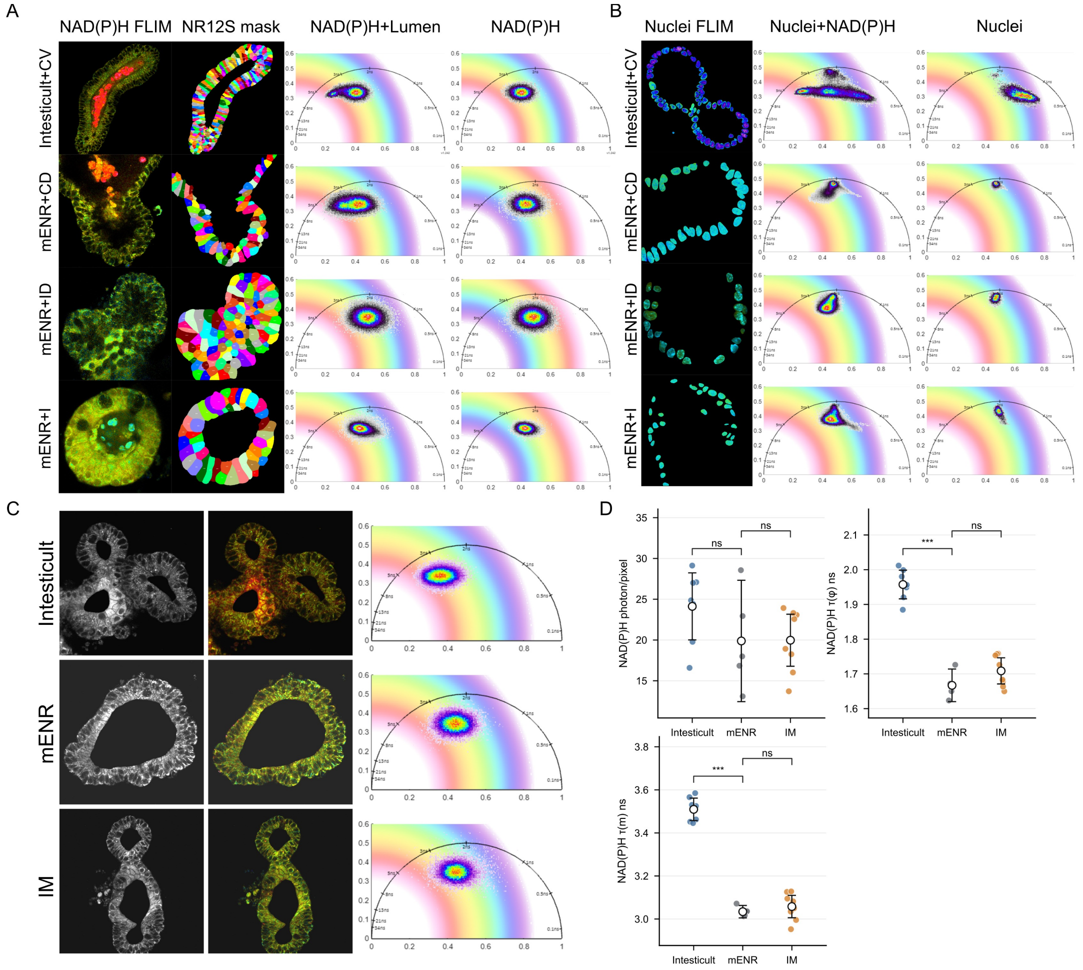


**Supplementary Figure S6. Phasor-based analysis of cellular and nuclear NAD(P)H signals in intestinal organoids under different differentiation and imaging conditions.**

**A**: Representative single-cell analysis of cellular NAD(P)H in intestinal organoids cultured under four differentiation conditions: IntestiCult + CV, mENR + CD, mENR + ID, and mENR + I. Cell masks were generated using Cellpose from NR12S membrane staining. Images are shown, from left to right, as NAD(P)H FLIM images, NR12S-based Cellpose masks, phasor plots containing cellular NAD(P)H and lumen-associated signals, and phasor plots displaying the selected cellular NAD(P)H signal. **B**: Representative single-cell analysis of nuclear fluorescence signals in intestinal organoids cultured under the four differentiation conditions. Images are shown, from left to right, as FLIM images, phasor plots containing both nuclear and cellular NAD(P)H signals, and phasor plots displaying the selected nuclear signal. **C**: Representative NAD(P)H intensity images, FLIM images, and corresponding phasor plots showing the effect of culture and imaging media on organoid NAD(P)H fluorescence lifetime. Organoids cultured in commercial IntestiCult or mENR were imaged directly in the same medium; the imaging-medium (IM; phenol-red-free, bicarbonate-free DMEM with 10 mM D-glucose, 2 mM GlutaMAX, 1 mM sodium pyruvate and 10 mM HEPES-Na, pH 7.2) group was transferred from mENR shortly before imaging. **D**: Quantification of cellular NAD(P)H intensity, τ(φ), and τ(m) in organoids imaged in IntestiCult medium, mENR medium, or IM.

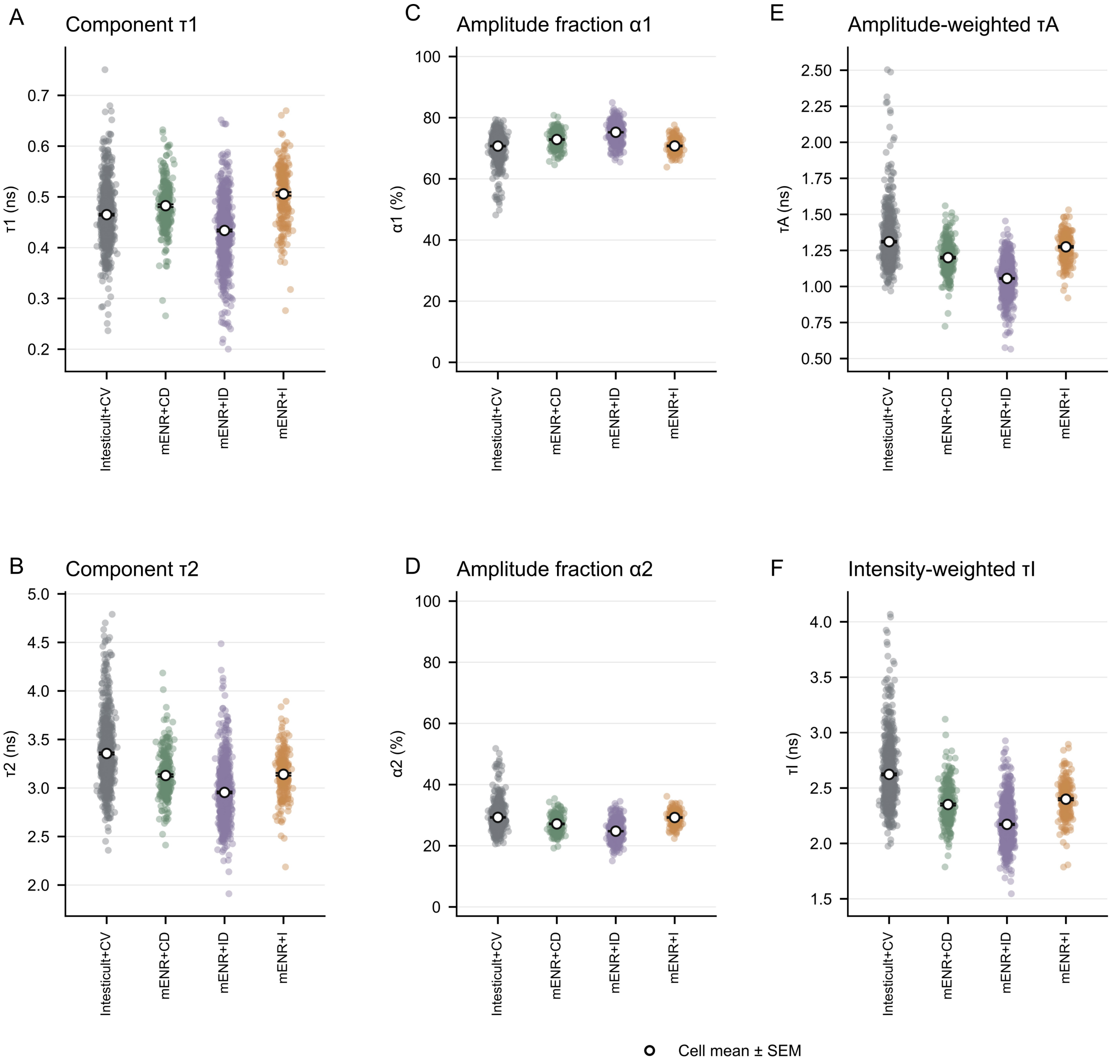


**Supplementary Figure S7. Calculated fitting decay parameters for NAD(P)H across four guided differentiation organoid culture conditions. A,B**: Short- and long-component lifetimes ($\tau_1$ and $\tau_2$). **C,D**: Corresponding amplitude fractions ($\alpha_1$ and $\alpha_2$). **E,F**: Amplitude-weighted (τA) and intensity-weighted (τI) lifetimes. Each point represents one independently fitted cell; white markers and error bars show the cell mean ± SEM. IntestiCult + CV, mENR + CD, mENR + ID and mENR + I include 883, 307, 853 and 255 cells from 6, 6, 10 and 5 images, respectively. All saved fitted values are included. Equal-image summaries and cell-level descriptive summaries are in Supplementary Table 11. Reconvolution and weighting are described in Methods, ‘Native fluorescence-decay fitting’.

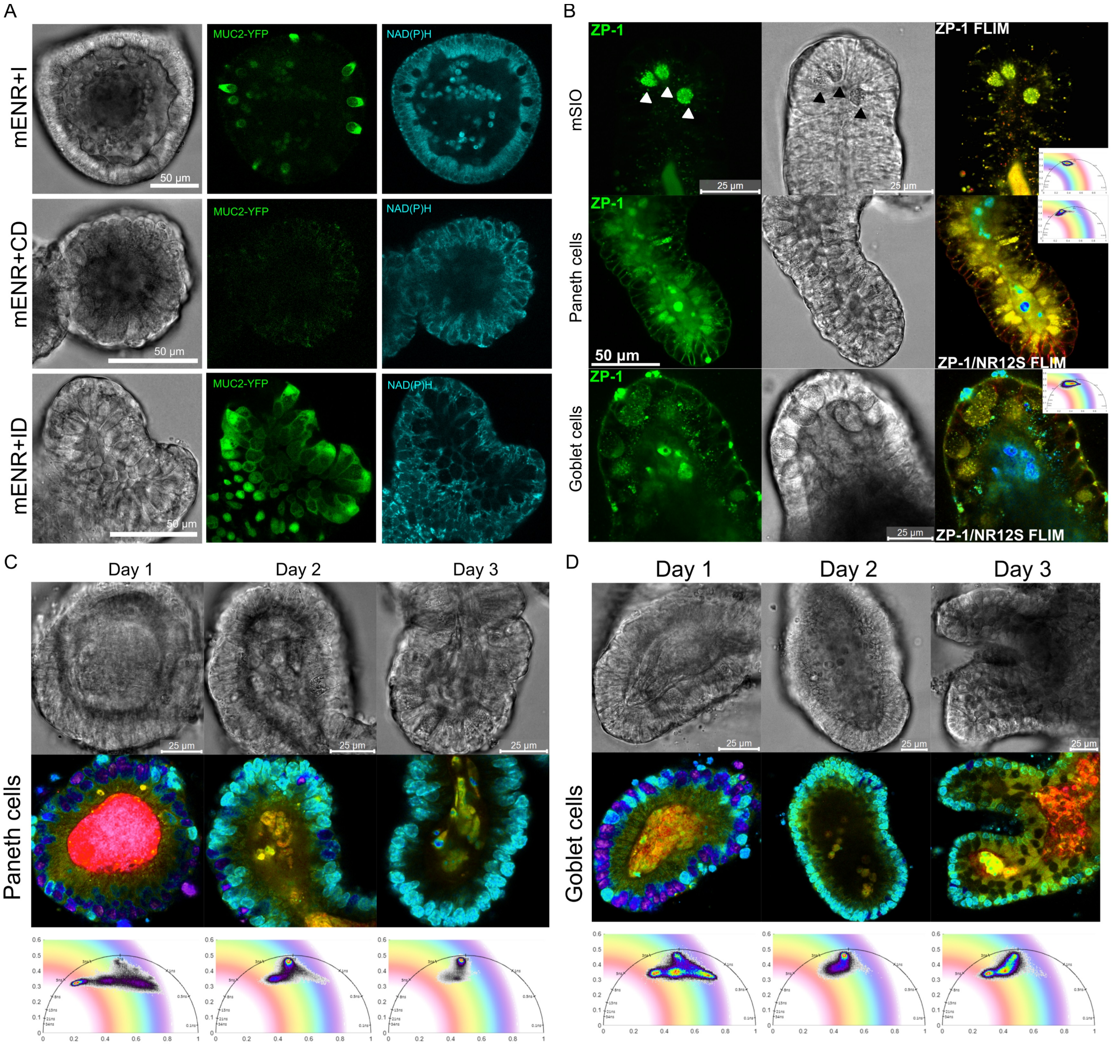


**Supplementary Figure S8. Fluorescent reporter and probe-based validation of differentiated intestinal organoid cell states.**

**A**: Representative bright-field, Muc2-YFP, and NAD(P)H images of Muc2-YFP intestinal organoids cultured under mENR + I, mENR + CD, and mENR + ID differentiation conditions. Under the mENR + I enterocyte differentiation condition, high NAD(P)H regions did not overlap with Muc2-YFP-positive cells. Under the mENR + CD Paneth cell differentiation condition, Muc2-YFP signal was absent. Under the mENR + ID goblet cell differentiation condition, the number of Muc2-YFP-positive cells increased, consistent with goblet cell enrichment. **B**: Representative Zinpyr-1 (ZP-1) staining in normal mouse small intestinal organoids (mSIOs), Paneth cell-enriched organoids, and goblet cell-enriched organoids. Normal mSIOs were stained with ZP-1 alone, whereas Paneth cell- and goblet cell-enriched organoids were co-stained with ZP-1 and NR12S. Images are shown, from left to right, as ZP-1 or ZP-1/NR12S intensity images, transmission light images, and ZP-1 or ZP-1/NR12S FLIM images. **C-D**: Representative BrdU–Hoechst 33342/NAD(P)H FLIM images showing the progression of Paneth cell and Goblet cell differentiation from day 1 to day 3 after stem cell enrichment in IntestiCult + CV. Images are shown, from top to bottom, as transmission light images, BrdU–Hoechst 33342/NAD(P)H-FLIM images, and corresponding phasor plots.

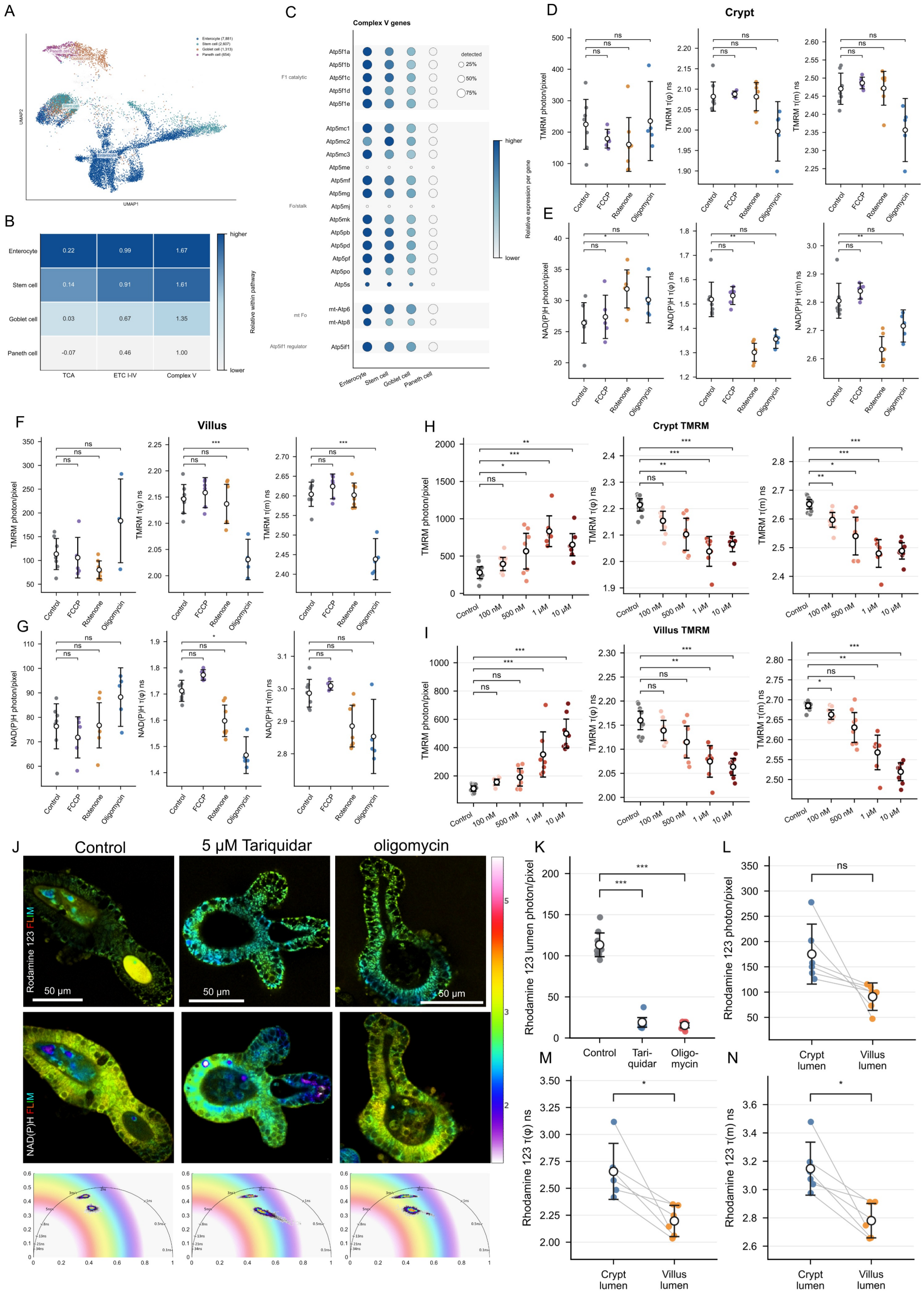


**Supplementary Figure S9. Oxidative metabolism-associated gene expression, mitochondrial perturbations and rhodamine 123 transport in intestinal organoids. A**: Uniform manifold approximation and projection (UMAP) of untreated mouse intestinal organoids cultured in EGF, Noggin and R-spondin (ENR), highlighting enterocytes (7,881 cells), stem cells (2,607), goblet cells (1,313) and Paneth cells (654); other populations are shown in grey. **B**: Expression-based scores for the tricarboxylic acid (TCA) cycle, electron transport chain complexes I–IV and nuclear-encoded ATP synthase (complex V) genes. Values are equally weighted means across five source-sample groups; colors indicate relative scores within each pathway. **C**: Expression of complex V-associated genes across the four cell types, grouped by ATP synthase component, with mitochondria encoded genes and the inhibitory regulator Atp5if1 displayed separately. Dot size indicates the percentage of cells with detectable transcripts; color indicates mean log-normalized expression scaled separately for each gene. **D–G**: TMRM intensity, τ(φ), and τ(m), in crypt **(D)** and villus **(F)** regions, with corresponding NAD(P)H measurements in crypt **(E)** and villus **(G)** regions, following 2 μM FCCP, 10 μM rotenone or 10 μM oligomycin treatment. TMRM was used at 10 nM; n = 7, 6, 7 and 5 images for control,

FCCP, rotenone and oligomycin, respectively. **H,I**: TMRM intensity, $\tau(\varphi)$ and $\tau(m)$ in crypt **(H)** and villus **(I)** regions following oligomycin treatment at 100 nM, 500 nM, 1 μM or 10 μM; n = 11, 9, 8, 7 and 8 images for control and increasing concentrations, corresponding to the dose series in Fig. 3D–H. **J**: Representative rhodamine 123 (5 μM) and NAD(P)H FLIM images under control conditions and following tariquidar (5 μM) and oligomycin (10 μM) treatment, with corresponding phasor distributions for both fluorophores. **K**: Luminal rhodamine 123 intensity under the conditions shown in J; n = 8, 9 and 10 images for control, tariquidar and oligomycin, respectively. **L–N**: Paired comparisons of rhodamine 123 intensity **(L)**, $\tau(\varphi)$ **(M)** and $\tau(m)$ **(N)** between crypt-adjacent and villus-adjacent luminal regions in control organoids; n = 6 paired images. Connecting lines indicate regions measured within the same image.

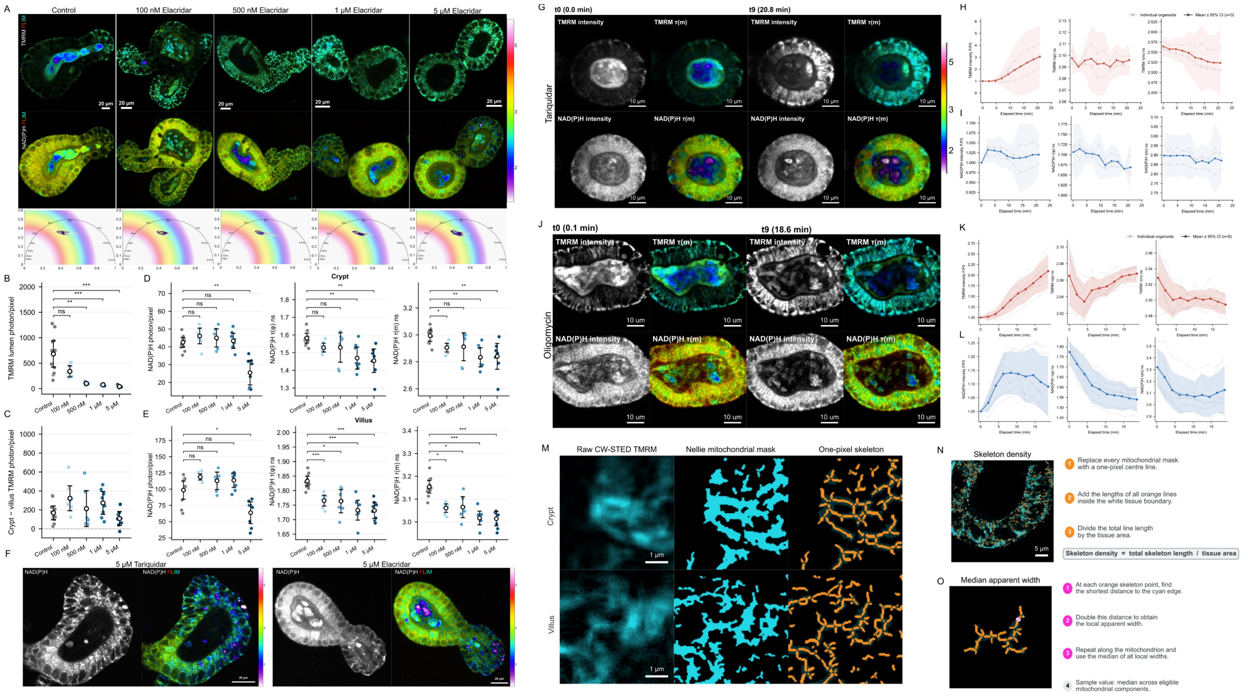


**Supplementary Figure S10. Elacridar responses, time-resolved TMRM–NAD(P)H-FLIM imaging and mitochondrial morphology analysis. A**: Representative TMRM and NAD(P)H-FLIM images and corresponding NAD(P)H phasor plots under control conditions and elacridar treatment conditions at 100 nM, 500 nM, 1 µM or 5 µM. TMRM was used at 10 nM. **B–E**: Quantification of luminal TMRM intensity **(B)**, paired crypt-minus-villus TMRM intensity differences **(C)**, and NAD(P)H intensity, τ(φ), and τ(m), in crypt **(D)** and villus **(E)** regions. Positive differences in C indicate greater crypt TMRM accumulation. For **B–E**, n = 11, 8, 7, 8 and 8 images for control and increasing elacridar concentrations, respectively; The control dataset is shared with the oligomycin dose series in Fig. 3 and Supplementary Fig. S9. **F**: Representative intensity and FLIM images showing bright, short-lifetime puncta in the NAD(P)H detection channel following 5 µM tariquidar or elacridar treatment. **G**: Representative villus-region TMRM and NAD(P)H intensity and τ(m) images at the initial (*t*0, 0.0 min) and final (*t*9, 20.8 min) acquisitions during the response to 500 nM tariquidar (Supplementary Video 1). **H,I**: Corresponding time courses of TMRM **(H)** and NAD(P)H **(I)** intensity, τ(φ) and τ(m); n = 3 sample trajectories. **J**: Representative intensity and τ(m) images at *t*0 (0.1 min) and *t*9 (18.6 min) during the response to 10 µM oligomycin (Supplementary Video 2). **K,L**: Corresponding TMRM **(K)** and NAD(P)H **(L)** time lapse measurements; n = 6 sample trajectories. Time-course intensities were normalized to the initial value within each sample and channel ($F/F_0$). Thin grey lines show individual trajectories, colored lines indicate means and shaded bands indicate 95% confidence intervals. **M**: Representative crypt and villus CW-STED TMRM images, Nellie-derived mitochondrial masks and one-pixel skeletons. Cyan denotes TMRM-positive structures and orange denotes skeletons. **N**: Skeleton-density analysis: total calibrated skeleton length within the manually delineated tissue boundary was divided by tissue area, excluding the lumen. **O**: Apparent-width analysis: local width was calculated as twice the Euclidean distance from each skeleton pixel to the mask background. Local widths were summarized by their median within each mitochondrial component, followed by the median across eligible components within each field.

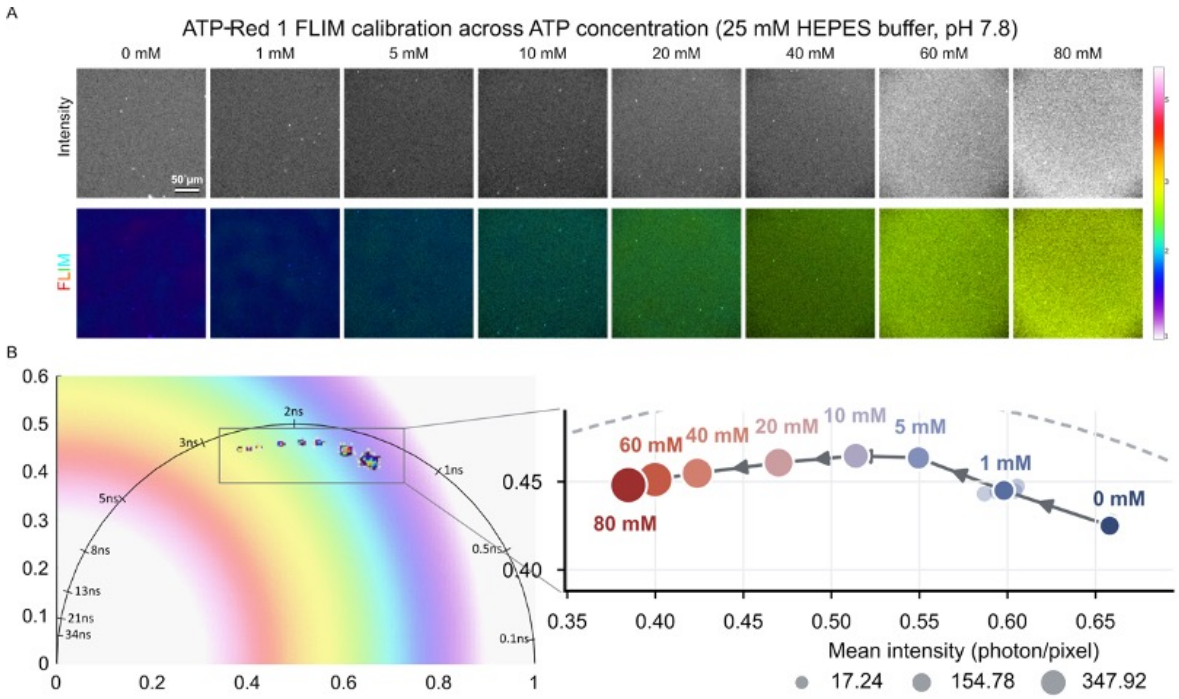


**Supplementary Figure S11. ATP-Red 1 fluorescence intensity and lifetime across the ATP calibration series. A**, Representative intensity and FLIM images of 5 µM ATP-Red 1 across ATP concentrations of 0–80 mM in 25 mM HEPES-Na buffer, pH 7.8. Intensity images were displayed using logarithmic rendering in GSLab. Scale bar, 50 µm. **B**, Corresponding phasor distributions and an enlarged view of the boxed region. Marker area encodes mean fluorescence intensity, and arrows indicate increasing ATP concentration. Five acquisition images were analyzed per concentration. Phasor trajectories are descriptive; solution calibration was not used to assign absolute intracellular ATP concentrations.

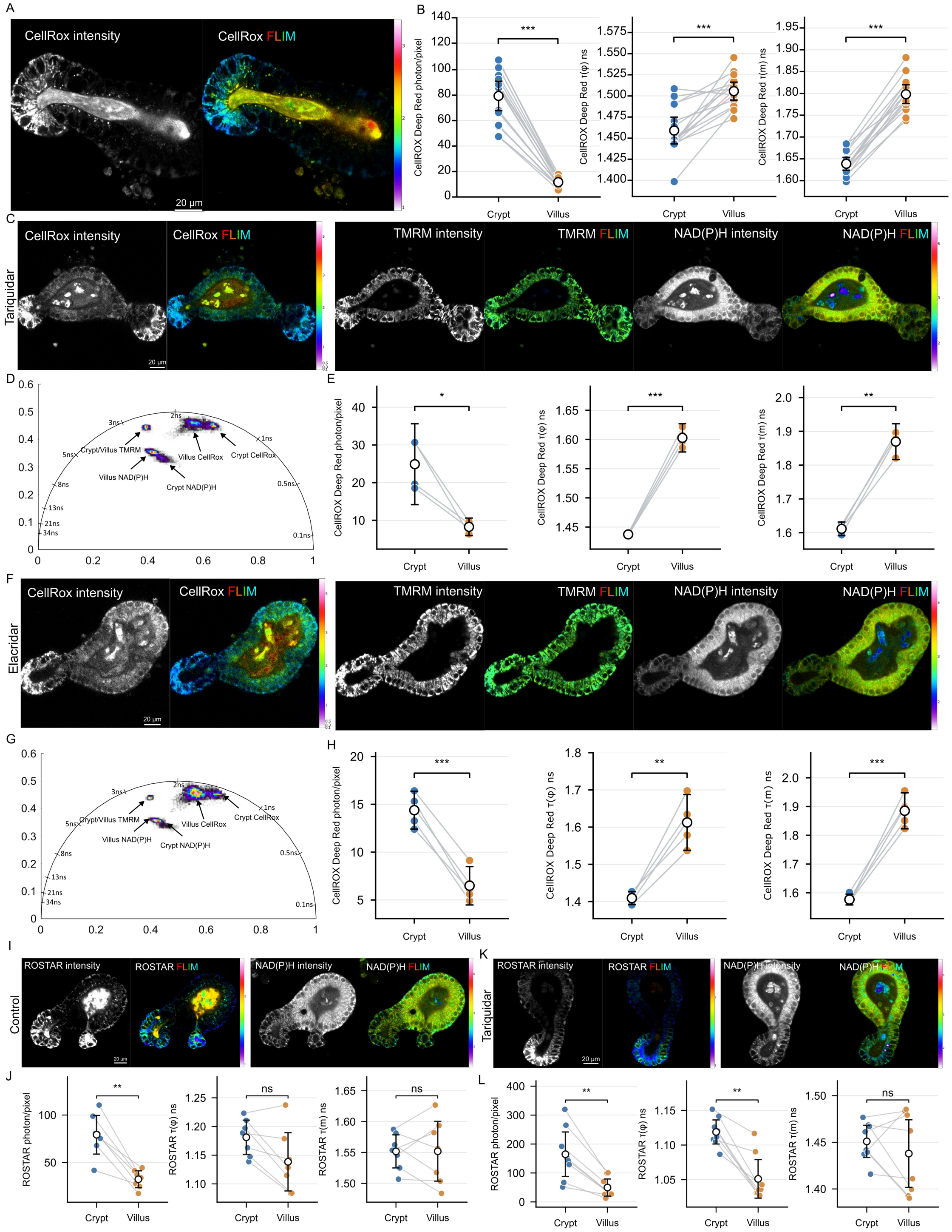


**Supplementary Figure S12. Spatial variability in ROS production observed with CellROX Deep Red and ROSstar 550 staining in live mouse intestinal organoids. A**: Representative CellROX Deep Red intensity and FLIM images under control conditions. **B**: Paired crypt–villus-like comparisons of CellROX Deep Red intensity, τ(φ), and τ(m); n = 15. **C**: Representative CellROX Deep Red, TMRM and NAD(P)H intensity and FLIM images following tariquidar treatment (500 nM). **D**: Corresponding phasor plot identifying regional fluorescence distributions. **E**: CellROX Deep Red intensity and lifetimes following tariquidar treatment; n = 4. **F–H**: Corresponding images **(F)**, phasor plot **(G)** and CellROX Deep Red measurements **(H)** following elacridar treatment (500 nM); n = 5. **I,J**: Representative ROSstar 550 and NAD(P)H intensity and FLIM images **(I)** and paired regional ROSstar 550 measurements **(J)** under control conditions; n = 7. **K,L**: Corresponding images **(K)** and ROSstar 550 measurements **(L)** following tariquidar treatment (500 nM); n = 8.

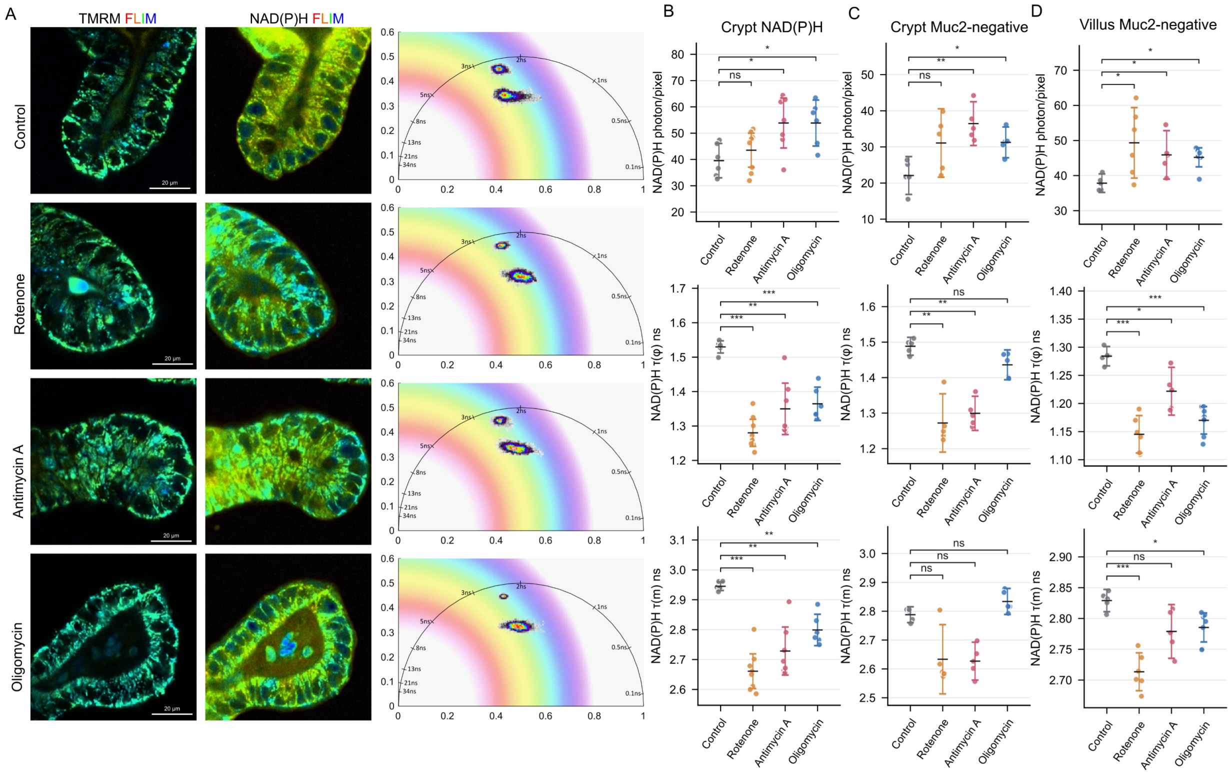


**Supplementary Figure S13. Regional NAD(P)H responses to the inhibition of electron transport and ATP synthesis in non-transfected wild-type and Muc2-YFP organoids. A,B**: Representative TMRM and NAD(P)H-FLIM images and corresponding phasor plots of wild-type intestinal organoid crypts **(A)**, and NAD(P)H intensity, τ(φ), τ(m), within TMRM-positive mitochondrial regions **(B)**, under control conditions or following rotenone, antimycin A or oligomycin treatment. **C,D**: Corresponding NAD(P)H measurements within TMRM-positive mitochondrial regions of Muc2-YFP-negative crypt **(C)** and villus **(D)** areas. Reporter-negative regions were defined as the complement of Cellpose-derived Muc2-YFP-positive cellular masks, excluding the reporter-positive populations analysed in Fig. 5 H,J and retaining mixed reporter-negative populations. Mitochondrial masks were generated separately for each image and treatment condition and applied to the corresponding NAD(P)H pixels.

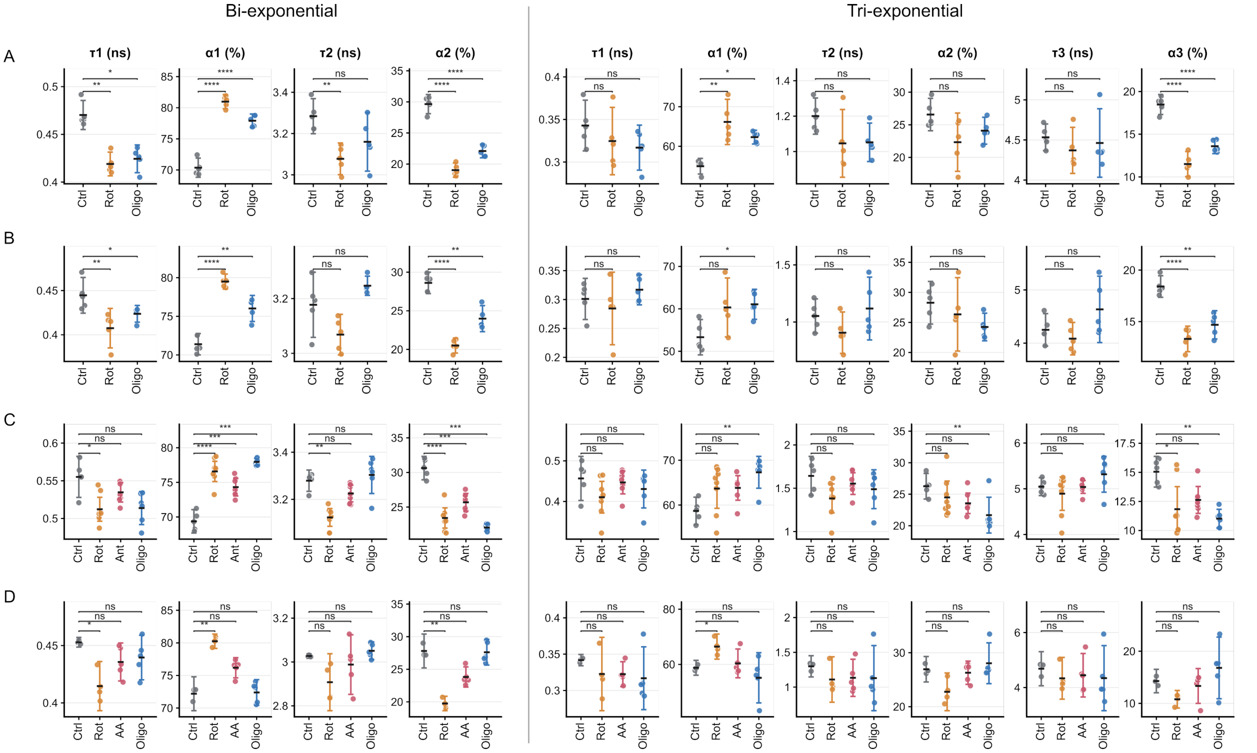


**Supplementary Figure S14. Bi- and tri-exponential analysis of cell-associated NAD(P)H decays across intestinal epithelial cell populations (Figure 5B, D, F, H)**

**A–D**: Component lifetimes ($\tau_i$, ns) and relative pre-exponential amplitudes ($\alpha_i$, %) for the datasets corresponding to Figures 5**B**, **D**, **F** and **H**, respectively. Bi-exponential fits are shown on the left and tri-exponential fits on the right. Stem-enriched and enterocyte measurements used TMRM-derived mitochondrial ROIs; Paneth-associated measurements used Muc2-YFP-positive cellular ROIs. Points represent independently fitted image ROIs; summaries show means and 95% confidence intervals. Existing treatment-versus-control comparisons and exact P values are presented in Supplementary Table 15. Ctrl, control; Rot, rotenone; Ant/AA, antimycin A; Oligo, oligomycin. Full-decay reconvolution and amplitude definitions are described in Methods, ‘Native fluorescence-decay fitting’.

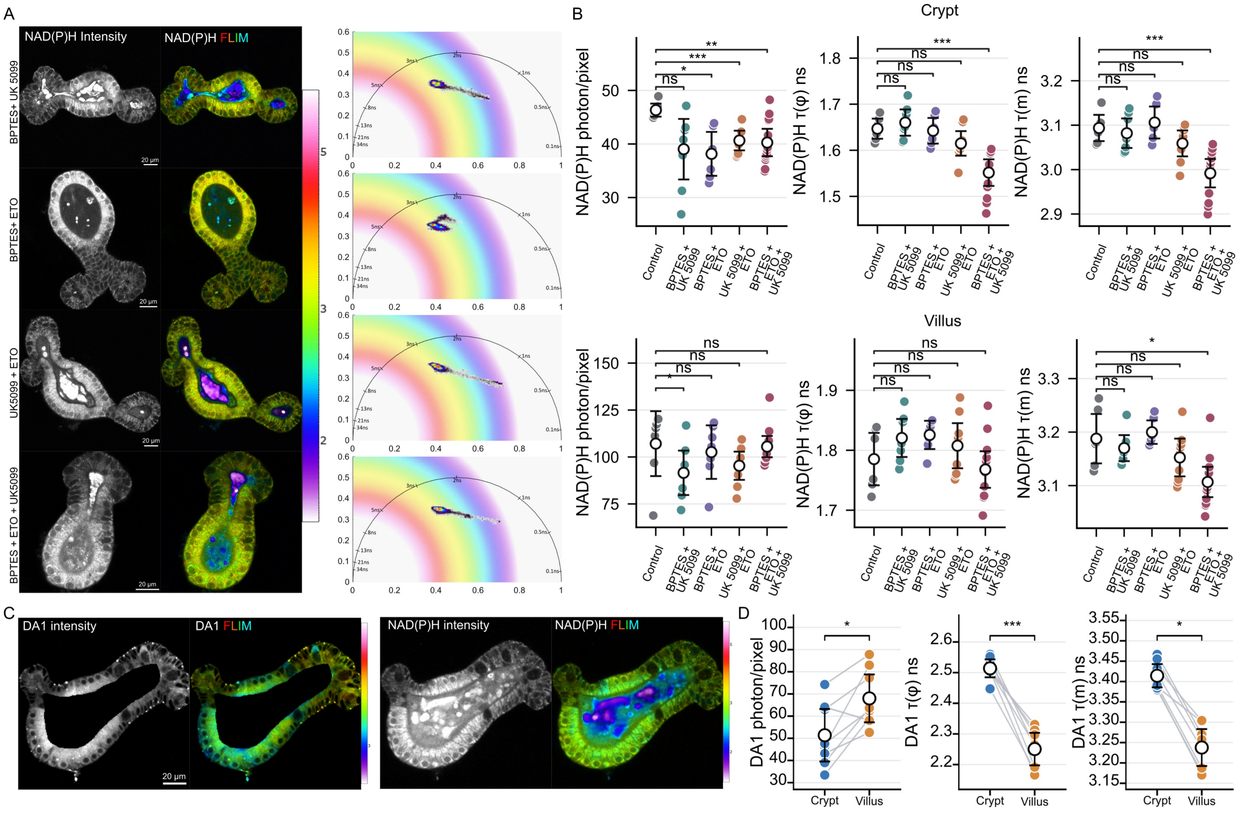

**Supplementary Figure S15. NAD(P)H responses to combined metabolic inhibition and spatial variation in lipids visualized with BDP-DA1 dye. A,B**: Representative NAD(P)H intensity and FLIM images with corresponding phasor plots for the indicated treatments **(A)**, and quantification of NAD(P)H intensity, phase lifetime, τ(φ), and modulation lifetime, τ(m), in crypt and villus regions **(B)**. Quantitative comparisons include control, BPTES + UK-5099, BPTES + etomoxir, UK-5099 + etomoxir and combined BPTES + UK-5099 + etomoxir treatment; n = 7 control, 8 BPTES + UK-5099, 7 BPTES + etomoxir, 9 UK-5099 + etomoxir and 13 triple-treatment images per region. **C,D**: Representative images of staining with BDP-DA1 and NAD(P)H intensity and FLIM images **(C)**, and paired comparisons of BDP-DA1 intensity, τ(φ) and τ(m) between crypt and villus regions **(D)**. Lines connect matched crypt and villus measurements. BDP-DA1 was excited at 514 nm and detected at 529–581 nm; n = 8 paired images.
